# Unselfish meiotic drive maintains heterozygosity in a parthenogenetic ant

**DOI:** 10.1101/2024.02.09.579553

**Authors:** Kip D. Lacy, Taylor Hart, Daniel J.C. Kronauer

**Affiliations:** Laboratory of Social Evolution and Behavior, The Rockefeller University, New York, NY, USA; Howard Hughes Medical Institute, New York, NY, USA

## Abstract

According to Mendel’s second law, chromosomes segregate randomly in meiosis. Non-random segregation is primarily known for cases of selfish meiotic drive in females, in which particular alleles bias their own transmission into the oocyte^1,2^. Here, we report a rare example of unselfish meiotic drive for crossover inheritance in the clonal raider ant, *Ooceraea biroi*. This species produces diploid offspring parthenogenetically via fusion of two haploid nuclei from the same meiosis^3^. This process should cause rapid genotypic degeneration due to loss of heterozygosity, which results if crossover recombination is followed by random (Mendelian) segregation of chromosomes^4,5^. However, by comparing whole genomes of mothers and daughters, we show that loss of heterozygosity is exceedingly rare, raising the possibility that crossovers are infrequent or absent in *O. biroi* meiosis. Using a combination of cytology and whole genome sequencing, we show that crossover recombination is, in fact, common, but that loss of heterozygosity is avoided because crossover products are faithfully co-inherited. This results from a programmed violation of Mendel’s law of segregation, such that crossover products segregate together rather than randomly. This discovery highlights an extreme example of cellular “memory” of crossovers, which could be a common yet cryptic feature of chromosomal segregation.

## Introduction

Meiosis and sex are ancient, predating the last common ancestor of eukaryotes^6^. Nevertheless, animal lineages that reproduce via parthenogenesis (development from eggs without fertilization by sperm) evolve sporadically via mutations that perturb meiosis. The underlying alterations to meiosis are rarely understood, but might provide insights into the evolution of asexuality and the fundamental mechanisms of meiosis.

Parthenogenetic lineages face two major challenges. First, they must restore euploidy without sexual fertilization, and second, they must produce offspring with viable genotypes. This often requires maintenance of heterozygosity to avoid recessive lethality, preserve genetic diversity, and transmit genotypes compatible with female development^5,7–9^. The many known parthenogenetic lineages employ a variety of cytological mechanisms for ploidy restoration^4,10^, but how they maintain heterozygosity is usually unknown.

The clonal raider ant, *Ooceraea biroi*, has satisfied these dual challenges. In this species, diploid offspring are formed via the fusion of two haploid pronuclei formed from the same meiosis following canonical meiotic divisions. Specifically, the second polar pronucleus fuses with one of the first polar pronuclei^3^. This mode of parthenogenesis, known as “automixis with central fusion,” solves the problem of ploidy restoration, but creates a problem for heterozygosity maintenance distal to crossovers (Figure 1a).

**Figure 1.**
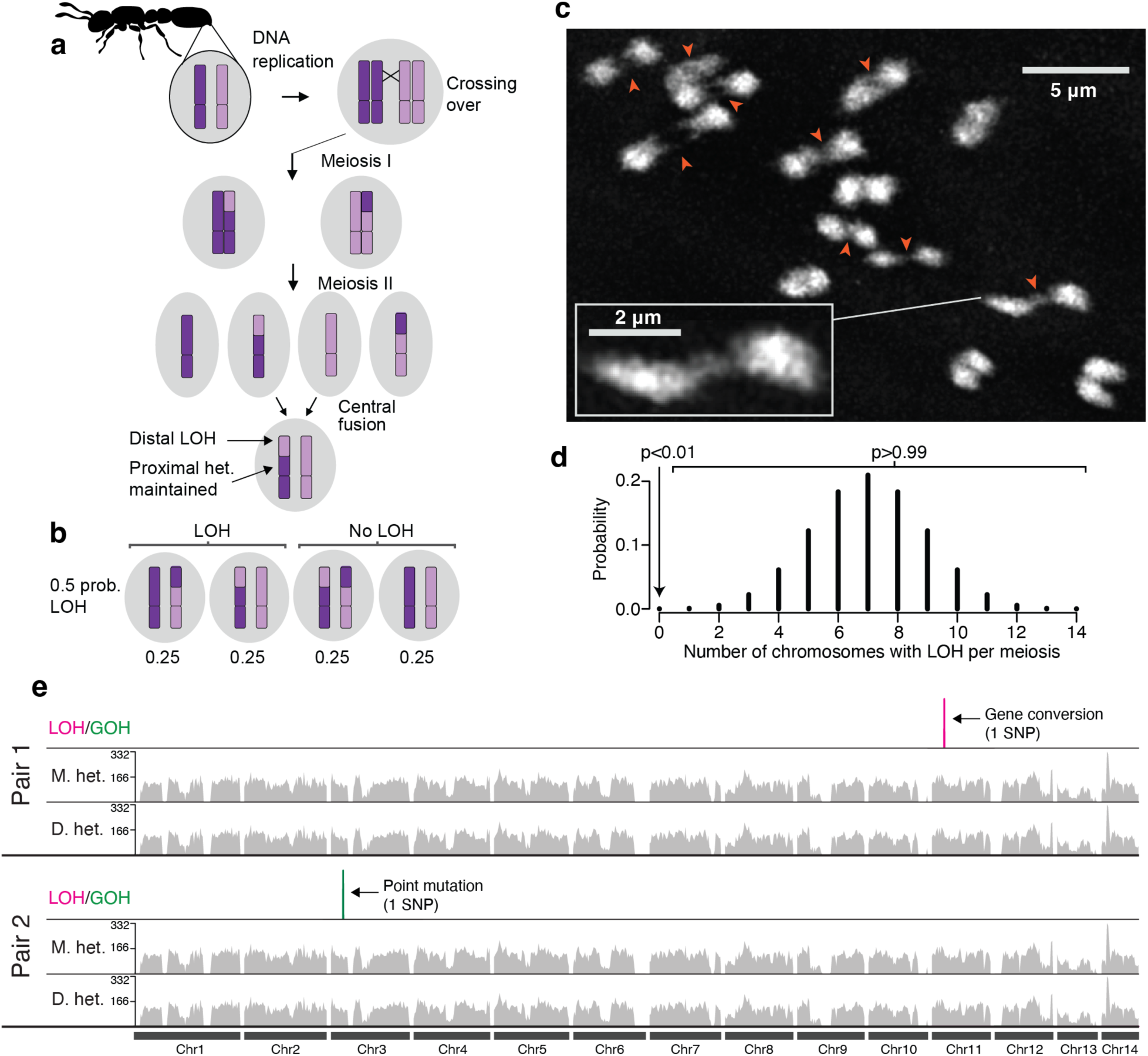
Loss of heterozygosity should be frequent but is rare in *O. biroi*. **a,** Schematic of *O. biroi* parthenogenesis, depicting the fusion of central haploid pronuclei to form the diploid zygote and the expected loss of heterozygosity distal to crossovers. Dark and light purple coloring on chromosomes indicates allelic identity. **b,** Depiction of all possible genomic compositions of diploid zygotes following segregation of a crossover and central fusion, along with their expected probabilities. **c,** Maximum projection of DAPI-stained *O. biroi* chromosomes at metaphase I, with pairs of homologs connected by chiasmata. Red arrows indicate clearly visible chiasmata. Pairs of homologs that lack red arrows are too close to each other to distinguish chiasmata clearly. **d,** Probability mass function of a binomial model describing how many chromosomes are expected to incur a loss of heterozygosity per meiosis, with one crossover on each of *O. biroi*’s 14 homologous chromosomes. Shown above are the probabilities that loss of heterozygosity occurs on no chromosomes or on one or more chromosomes. **e,** Karyoplot depicting all SNPs that underwent loss-of-heterozygosity (magenta) or gain-of-heterozygosity (green) among the sequenced genomes from two mother-daughter pairs. The expected segmental losses of heterozygosity were notably absent. For each individual, grey histograms show the number of sites heterozygous in 250kb windows.

Crossovers provide the physical tension that ensures accurate chromosome segregation in meiosis. Accordingly, most species have at least one crossover per chromosome (crossover assurance). Following a crossover, chromosomes are thought to segregate randomly (Mendel’s second law). In central fusion parthenogenesis, there are four possible inheritance outcomes following a crossover, and random segregation means that they are expected to occur with equal probability (Figure 1b). Two of these outcomes lead to segmental loss of heterozygosity due to the inheritance of one crossover product and one non-crossover product. Thus, we expect half of all crossovers to produce a segmental loss of heterozygosity. If crossovers occur on each of *O. biroi*’s 14 homologous chromosomes^11,12^, then loss of heterozygosity should occur commonly. However, genetic studies suggest that little, if any, heterozygosity is lost each generation^3,13^. How heterozygosity is maintained has remained a mystery.

## Results

### Loss of heterozygosity should be common, but it is rare

One possibility is that *O. biroi* meiosis lacks crossovers. Crossovers are required in canonical meiosis^14^, but are absent from male or female meiosis in many insect species (sex-limited achiasmy)^15^. This is shown using chromosome squashes, which make physical crossover structures (chiasmata) visible as DNA connections between paired homologous chromosomes^16,17^. Achiasmate meiosis lacks such connections^18^. To determine whether *O. biroi* meiosis lacks crossovers, we performed chromosome squashes during metaphase I. We found that paired homologous chromosomes were linked by positive DAPI staining, indicative of chiasmata (Figure 1c, Supplementary Figure 1, and Supplementary Video 1). This suggests that crossovers occur normally in *O. biroi* meiosis, which means losses of heterozygosity should be frequent.

How frequent should losses of heterozygosity be, exactly? To answer this question, we used the binomial distribution, which is a good fit because 1) assuming Mendelian segregation, each crossover has a fixed probability of 0.5 of leading to segmental loss of heterozygosity, and 2) homologous chromosomes comprise a set of independent trials. In this way, we obtained the probability distribution of the number of chromosomes that undergo at least one loss of heterozygosity each meiosis (Figure 1d). This distribution showed that loss of heterozygosity should typically occur on multiple chromosomes, with a >99% chance of occurring on at least one chromosome.

Previous studies reported that loss of heterozygosity was rare, but relied on sparse genetic markers and compared individuals of unknown pedigree^3,13^. Tracking genotypes across generations in clonal colonies is challenging, so we created a transgenic clonal line with broad expression of the fluorescent marker protein dsRed (see Methods, Supplementary Figures 2 and 3, and Supplementary Table 1). To examine the amount of heterozygosity lost during *O. biroi* meiosis in detail, we then sequenced whole genomes of two mother-daughter pairs (descriptions and metadata for all genomes sequenced in this study are in Supplementary Table 2). No segmental losses of heterozygosity occurred in either mother-daughter pair (Figure 1e; Supplementary Table 3). Moreover, no segmental losses of heterozygosity occurred since the common ancestor of the mothers of the two pairs, even though these two genomes were separated by at least four but fewer than 24 meioses. The only changes in heterozygosity found between any of these individuals spanned short genomic tracts, including gains of heterozygosity via point mutations and small losses of heterozygosity due to gene conversion, which occurs at the site of both crossover and noncrossover recombination and affects regions typically less than 1500bp in length^19,20^ (Supplementary Table 4). One mother-daughter pair suffered loss of heterozygosity at a single SNP, probably due to gene conversion; the other pair gained heterozygosity at a single SNP due to a point mutation. Between the two mothers, there were twelve short losses of heterozygosity consistent with gene conversion and two gains of heterozygosity due to point mutations. These data reveal that, paradoxically, segmental loss of heterozygosity is extremely rare.

### Crossovers occur every meiosis without loss of heterozygosity

Taken together, these results imply that *O. biroi* employs a previously unknown mechanism to maintain heterozygosity. One possibility is that crossovers do not occur, and the putative chiasmata in chromosome squashes result from another kind of DNA entanglement. To explore this possibility, we looked for genetic evidence of crossovers, which can be challenging to study because, in diploid asexual lineages, they can only be identified using loss of heterozygosity as a proxy (Figure 2a). However, sequencing haploid genomes allows identification of inherited crossovers regardless of whether heterozygosity was lost. Fortunately, *O. biroi* clonal lines occasionally produce vestigial haploid males, presumably via meioses not followed by central fusion. We sequenced whole genomes of several haploid males and diploid females from two clonal lines (Line A, colony C16; Line B, colony STC6). This revealed hundreds of crossovers among haploid genomes, which contrasted with only a few crossovers inferred via loss of heterozygosity among diploid genomes (Figure 2b,c, Supplementary Data 1; haploids – Line A: 346 crossovers, Line B: 418; diploids – Line A: 8, Line B: 3). This effect was specific to crossover recombination, as haploids and diploids had roughly equal numbers of gene-conversion-sized events (haploids – Line A: 47 gene conversions, Line B: 32; diploids – Line A: 49, Line B: 51). Many more crossovers are thus evident by comparing haploid genomes than by observing losses of heterozygosity among diploid genomes.

**Figure 2.**
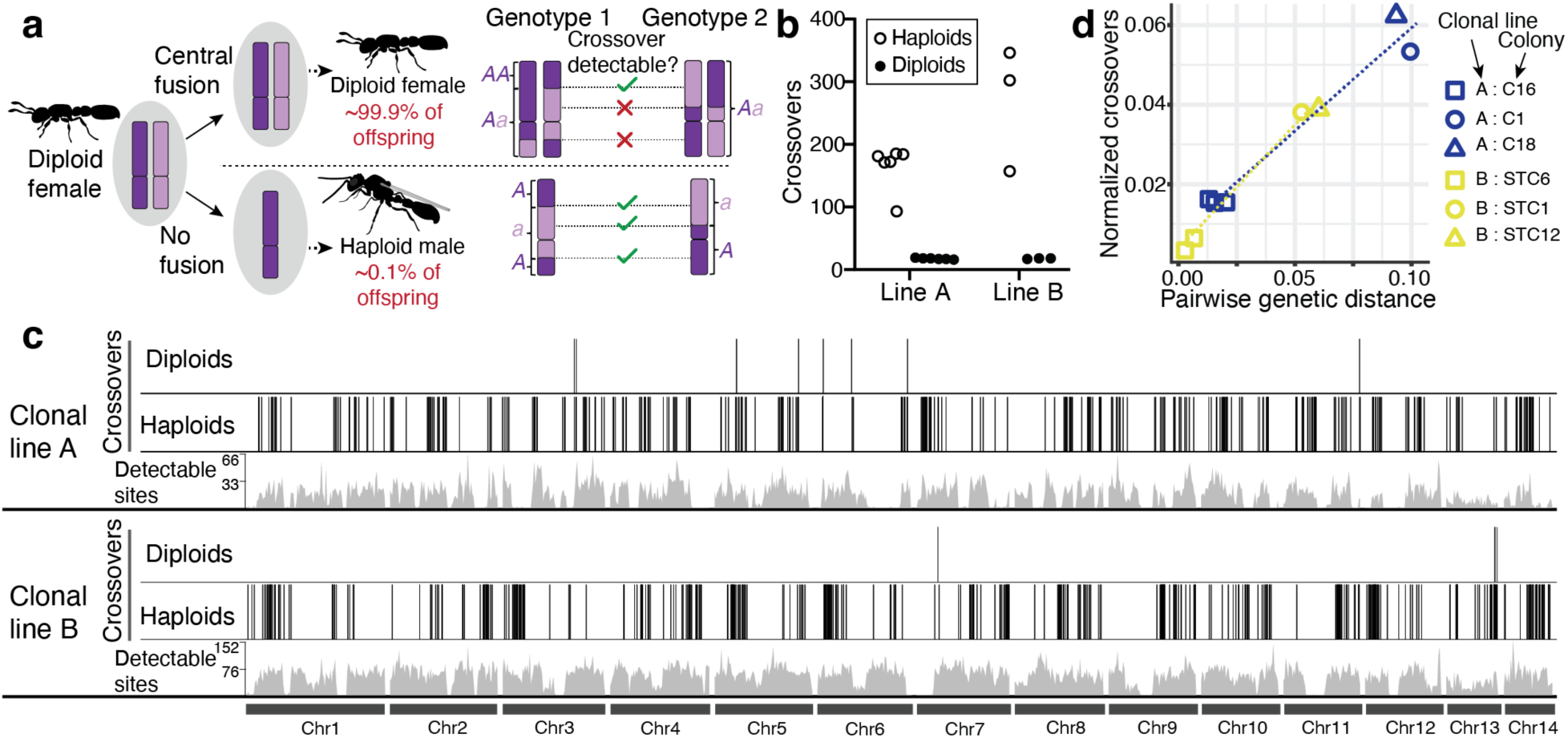
Haploid male genomes indicate that crossover recombination is common. **a,** Haploid males occur rarely in *O. biroi*, but can be used to detect crossover recombination. Light and dark purple coloring on chromosomes indicates allelic identity, and purple lettering next to chromosomes represents genotypes inferred from whole-genome sequencing. A comparison of example genotypes between two individuals illustrates that, using short-read sequencing, more crossovers are detectable between haploid genomes than between diploid genomes. This is because crossover recombination is directly observed in haploid males and inferred from segmental losses of heterozygosity in diploid females. Therefore, crossovers not followed by loss of heterozygosity cannot be detected among diploid genomes. **b,** The numbers of crossovers detected among all pairwise comparisons of haploid male and diploid female genomes. We sequenced four haploids and four diploids from a single stock of clonal line A, yielding six pairwise comparisons, and three haploids and three diploids from a stock of clonal line B, yielding three pairwise comparisons. **c,** Karyoplot illustrating all detected crossovers (dark vertical bars) and the number of sites ancestrally heterozygous in 200kb windows (“Detectable Sites”; grey histograms). Closely spaced vertical bars (crossovers) appear as thick black blocks. Many more crossovers are detected among haploids than among diploids. **d,** Jittered scatterplot depicting the correlation of crossover recombination with genetic distance in two *O. biroi* clonal lines. Within each clonal line, each point represents the number of crossovers observed between each male and a randomly selected male from the reference colony (C16 for clonal line A; STC6 for clonal line B). To examine data spanning greater genetic distances we sequenced one male each from two additional stock colonies from each clonal line. Pairwise genetic distances were calculated between diploid females from the respective colonies. Dotted lines depict linear models. The positive slope and near-zero y-intercepts indicate that, rather than excess crossovers being a quirk of haploid male production, crossovers accumulate gradually over generations of diploid-producing parthenogenesis.

*Ooceraea biroi* females do not mate in the lab, and males are evolutionary dead-ends in that they do not produce offspring. Therefore, since the common ancestor of the studied males, all meioses were followed by central fusion, until each male resulted from a final meiosis not followed by central fusion. This implies two possible alternative explanations of the dramatic differences in observed vs. inferred crossover events between males and females, respectively (Supplementary Figure 5). First, the meioses that produce haploid offspring might not only lack the subsequent central fusion, but also differ from typical meioses in other ways. Specifically, our data could be explained if typical meioses lacked crossovers, but the haploid-producing meioses featured excessive recombination. Alternatively, the crossovers observed among haploid male genomes might have accumulated over many preceding generations of females. In that case, crossovers should also be present but “hidden” in diploid female genomes because no heterozygosity was lost. To distinguish between these scenarios, we sequenced additional genomes from different colonies within each clonal line. This multi-colony dataset confirmed our initial results, also revealing many more crossovers among haploid than among diploid genomes (Supplementary Figure 6, Supplementary Data 2). We then analyzed how the number of crossovers among haploid males scaled with phylogenetic distance. If haploid-producing meioses featured excessive recombination, then we would expect non-zero y-intercepts, because closely related males would differ from each other by many recombination events. However, for both clonal lines, the y-intercepts were approximately zero (Figure 2d; Line A: 0.007; Line B: 0.004), implying that haploid males do not arise from meioses with exceptional numbers of crossovers. Moreover, the number of crossovers between males was tightly correlated with genetic distance (Line A: R^2^=0.94, p=0.007; Line B: R^2^=0.99, p=0.002), revealing that crossovers had accumulated over many generations and thus must occur regularly without loss of heterozygosity in *O. biroi* meiosis.

To confirm that crossovers occur without segmental loss of heterozygosity, we need to compare haplotypes from known pedigrees. Because haploid males occur too rarely to be used in pedigree studies, we used linked-read sequencing to obtain haplotype-resolved diploid genomes of five mother-daughter pairs (one meiosis each) and four sister-sister pairs (two meioses each). For each individual, we sequenced the linked-read library to at least 50x coverage with Illumina, allowing us to phase 61% of the genome on average, with an average phased block N50 of 27 kb (Supplementary Figure 7 and Supplementary Table 2). By comparing pairs of phased genomes, we can detect crossovers in heterozygous regions that are reliably phased in both samples (55% of the genome on average) and for which at least one recombined chromatid has been inherited. Crossovers are not detectable if neither recombined chromatid has been inherited, which is expected for one in four crossovers (Figure 1b). We detected crossovers in all known-pedigree pairs, with an average of 2.38 crossovers per meiosis (Figure 3 and Supplementary Table 5; 31 crossovers sampled across the 13 meioses from known-pedigree pairs). Crossovers were distributed across all chromosomes except for the heteromorphic chromosome 13, for which pervasive structural variation severely limits crossover detection^21^. Adjusted for underestimation due to incomplete phasing and non-detection of uninherited crossovers (see Methods), these data suggest that, on average, 5.5 crossovers occur in each meiosis (0.4 crossovers per chromosome, 95% CI 0.3-0.49). This is likely an underestimate because recombination hotspots might fall within regions that have ancestrally lost heterozygosity, and non-crossover products might disproportionately be co-inherited. Regardless, our data show that crossovers occur in every meiosis.

**Figure 3.**
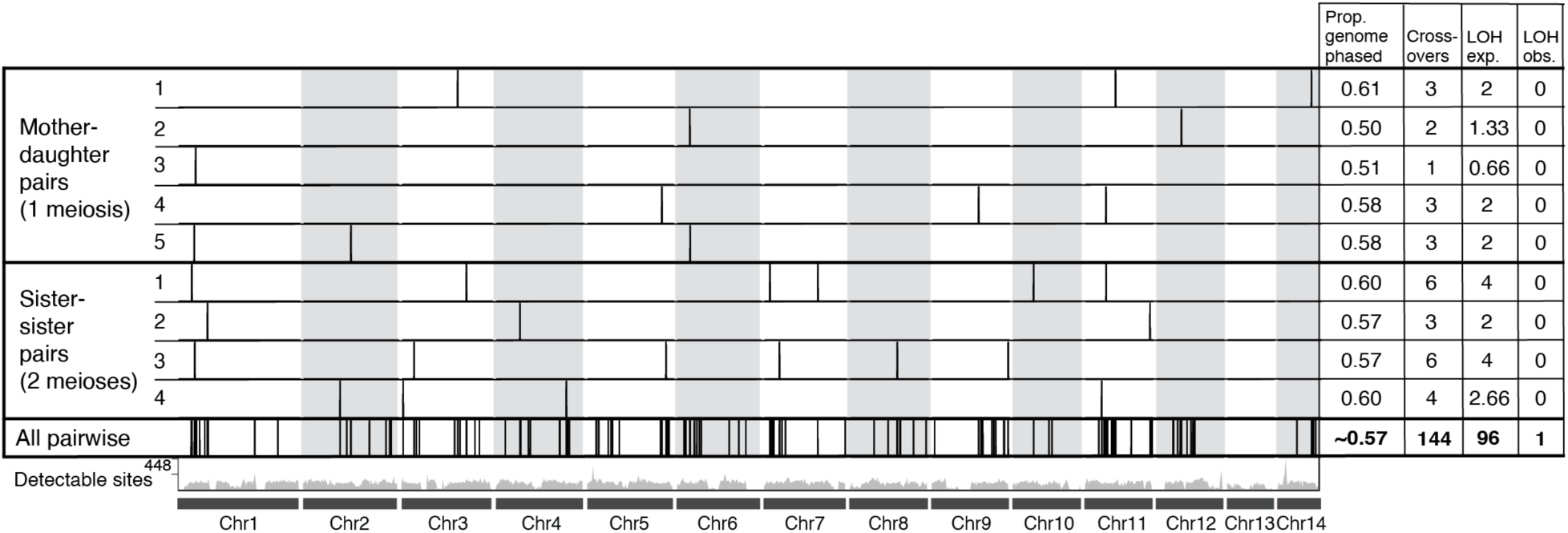
Crossovers occur every meiosis without loss of heterozygosity. Karyoplot depicting crossovers (tick marks) inferred between each pair of phased genomes from linked-read sequencing data and all pairwise comparisons (“All Pairwise”) between the different families of unknown pedigree. The numbers of sites at which crossovers are detectable per 200kb window are shown as a grey histogram (“Detectable Sites”). For each pair, we display to the right of the karyoplot the proportion of the genome phased in both samples (i.e., the proportion of the genome for which crossovers are detectable), the total number of crossovers detected, the number of losses of heterozygosity expected, and the number of losses of heterozygosity observed. Assuming Mendelian segregation, two-thirds of inherited crossovers should be associated with loss of heterozygosity, but surprisingly, only one of the observed crossovers was.

None of the 31 identified crossovers in this analysis were associated with segmental loss of heterozygosity, although 15 were associated with gene conversion (Figure 3 and Supplementary Table 5). This is surprising because, under Mendelian inheritance, we expect 2/3 of inherited crossovers to be associated with segmental loss of heterozygosity (Figure 1b). To investigate whether this pattern held for larger numbers of crossovers, we selected one individual from each pedigree, and performed all pairwise comparisons of their phased genomes (seven different pedigrees with 21 pairwise comparisons). Pairs of genomes in this analysis were separated by at least three but fewer than 25 meioses. This produced a total of 144 crossovers (including those that were detected in known-pedigree pairs). Of these, 143 crossovers occurred without segmental loss of heterozygosity, while one resulted in a >200kb loss of heterozygosity tract (Figure 3 and Supplementary Table 5, Supplementary Data 3 & 4). Thus, both crossover products were inherited for all but one detected crossover. This is a statistically significant departure from the expectation that segmental loss of heterozygosity should occur for 2/3 of inherited crossovers (Binomial Test, p<0.0001). The only other changes in heterozygosity were small (≤466 bp) losses of heterozygosity consistent with gene conversion (Supplementary Tables 4 & 5, Supplementary Data 3). This high-fidelity heterozygosity maintenance is showcased by sliding window analyses, which reveal remarkable stability of genome-wide heterozygosity levels (Supplementary Table 3). From a per-offspring perspective, loss of heterozygosity is also very rare; of the 15 offspring sampled (from both linked-read and short-read sequencing), none had a segmental loss of heterozygosity (95% CI: 0 – 0.2 for the true proportion of offspring that incur a segmental loss of heterozygosity).

### Programmed co-inheritance of crossover products maintains heterozygosity

These data cannot be explained via Mendelian inheritance of crossover products (Figure 4a). The underlying non-Mendelian mechanism must operate after homologous chromosomes separate (in anaphase I of meiosis), but before the offspring genotype is determined when central pronuclei fuse to form the diploid zygotic nucleus. In this period, chromosomes segregate during meiotic divisions. Under Mendelian inheritance, the segregation of one crossover product does not depend on the reciprocal crossover product. If this were the case in *O. biroi*, losses of heterozygosity would be the rule; even with our conservative empirical estimate of 0.4 crossovers per chromosome per meiosis, binomial modeling shows that, if segregation were Mendelian, 95% of offspring (95% CI 90%-98%) would bear a loss of heterozygosity on one or more chromosomes (Figure 4b). Therefore, chromosomal segregation must be non-random, such that if one crossover product is included in a central pronucleus, then the reciprocal crossover product is found in the other central pronucleus (Figure 4a). Such co-segregation of crossover products would ensure that inheritance of both (or neither) of the crossover products is the rule, whereas inheritance of one crossover product and one noncrossover product is a rare exception (Figure 4a,c).

**Figure 4.**
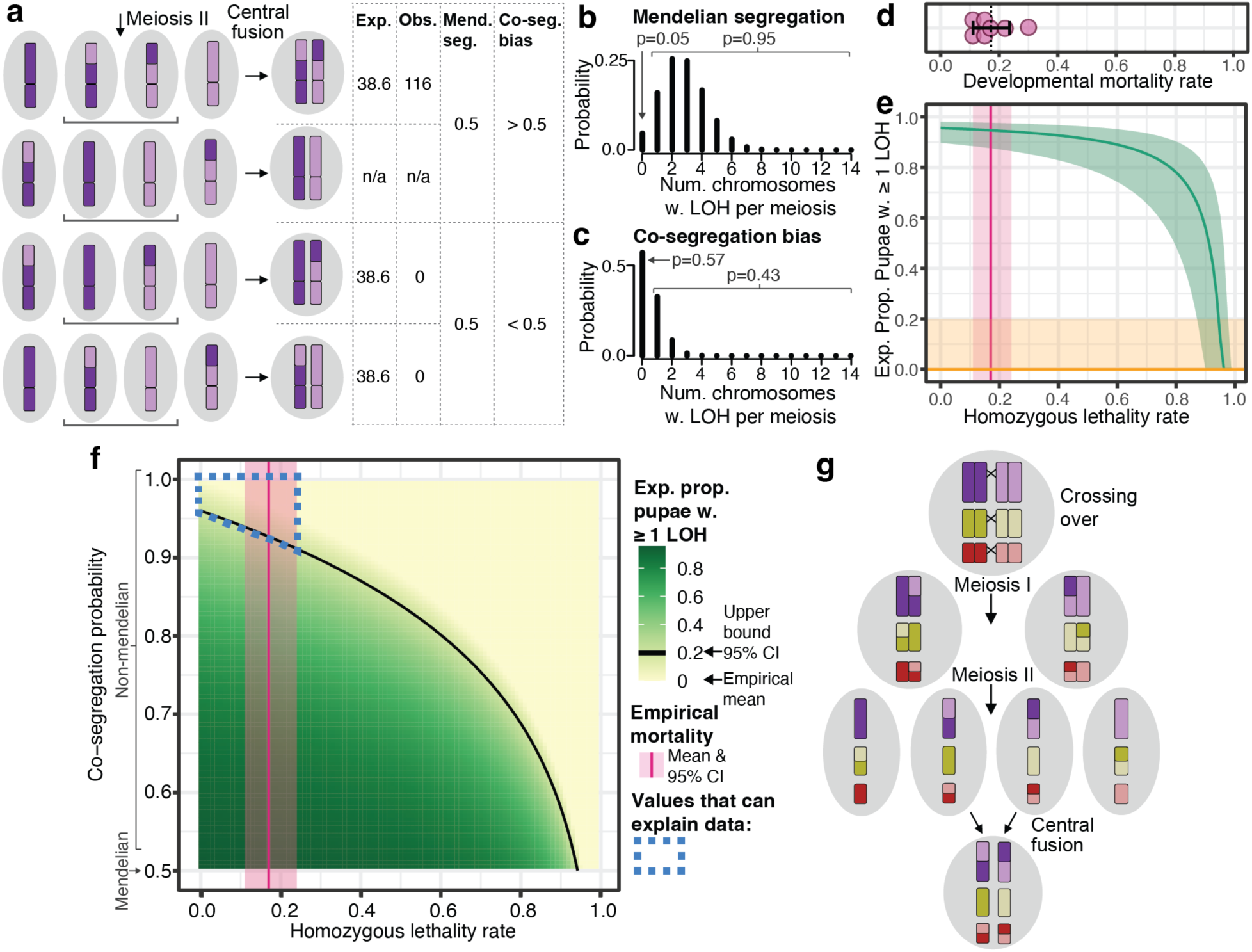
Non-Mendelian co-segregation of crossover products maintains heterozygosity. **a,** Cartoon depicting the four possible segregation patterns following a crossover. Shown are the stage following meiosis II, at which there are four haploid pronuclei, and the stage following central fusion, when the diploid zygote is formed. For each pattern, the expected and observed numbers of crossovers detected by linked-read sequencing are shown. Crossovers cannot be detected if neither crossover product was inherited, so the numbers for that outcome are listed as “n/a.” Under Mendelian segregation, the probability of loss of heterozygosity is 0.5, but such values are incompatible with our results. If crossover products co-segregate (“Co-segregation Bias”) such that they are always inherited together, heterozygosity would rarely be lost. **b,c,** Probability mass functions of binomial models depicting the number of chromosomes expected to incur a loss of heterozygosity per meiosis, assuming the empirical crossover rate of 5.5 crossovers per meiosis. The model in **b** assumes Mendelian segregation, whereas the model in **c** assumes a co-segregation probability of 0.9 (chosen because, at this probability, it is more likely for heterozygosity to be lost for zero chromosomes than for one or more chromosomes). Above the plots are the probabilities that losses of heterozygosity occur on zero chromosomes or on one or more chromosomes. **d,** Developmental mortality, shown as the proportion of individuals that died between the early egg stage and pupation in replicate colonies (magenta dots); mean (dotted line) and 95% CI (black bar). **e,** Theoretical model depicting the ability of homozygous lethality to maintain heterozygosity if segregation is Mendelian. Shown in green is the expected proportion of offspring with a loss of heterozygosity on one or more chromosomes (from the binomial model depicted in **b**) minus the homozygous lethality. Values based on the mean empirical crossover rate are shown as a dark line, and values based on the 95% CI are shown in green shading. The empirical developmental mortality rate (which is the hypothetical maximum homozygous lethality) is shown in magenta (mean depicted by dark line, 95% CI depicted by shading), revealing that at least 86% of offspring should bear a loss of heterozygosity. The empirical proportion of offspring with at least one loss of heterozygosity is shown in orange (mean depicted by dark line, 95% CI depicted by shading), revealing that developmental mortality rates would need to be at least 0.87 for homozygous lethality to produce proportions that fall within the observed range. Thus, homozygous lethality can have at most a small effect on heterozygosity maintenance. **f,** Heatmap depicting the proportion of offspring expected to bear a loss of heterozygosity on one or more chromosomes. “Reasonable” parameter combinations can be found above the black line, which depicts proportions of 0.2, the upper bound of our 95% CI for proportions of offspring that bear one or more LOH. The empirical developmental mortality rate (the hypothetical maximum homozygous lethality) is shown in magenta. Therefore, the only parameter combinations that can explain the degree of heterozygosity maintenance observed in this study are found within the blue dashed outline, revealing that a co-segregation probability greater than 0.91 occurs in *O. biroi*. **g,** Heterozygosity is maintained via a programmed violation of Mendelian segregation, whereby crossover products co-segregate such that either both or neither are inherited.

The only possible alternative to crossover co-segregation would be that losses of heterozygosity are lethal. Such “homozygous lethality” is thought to maintain heterozygosity in other parthenogenetic species, including the cape honeybee, which also reproduces via automixis with central fusion^22,23^. Because egg laying precedes meiotic divisions and central fusion in *O. biroi*^3^, homozygous lethality would occur after eggs are laid, but before the pupal stage, when we measured loss of heterozygosity. To measure developmental mortality, we used our transgenic marker line to track survival from the egg to the pupal stage in seven replicate colonies (Supplementary Figure 3). Mean mortality was 0.17 (95% CI 0.11-0.24) (Figure 4d). We then adjusted our binomial model, making the strict assumptions that 1) all developmental mortality results from loss of heterozygosity and 2) all losses of heterozygosity are lethal. This revealed that mortality rates of at least 0.87 would be required to produce proportions of offspring with loss of heterozygosity that fall within the 95% CI of our empirical estimate (Figure 4e). By contrast, our observed developmental mortality rate predicts expected proportions of offspring bearing loss of heterozygosity to be at least 0.86 under Mendelian segregation (Figure 4e). Therefore, even under our conservative estimate of crossover rates, homozygous lethality alone cannot maintain heterozygosity in *O. biroi*. Instead, crossover product co-segregation bias is required to explain our data.

To estimate the strength of this bias, we varied the probability of crossover product co-segregation as a parameter in our binomial model. Because we cannot rule out a role for homozygous lethality, we calculated the expected proportion of offspring with loss of heterozygosity for a range of homozygous lethality rates and co-segregation probabilities. Taking the upper bound of the developmental mortality rate as the hypothetical maximum homozygous lethality, we find that the crossover product co-segregation probability must be at least 0.91 for heterozygosity to be maintained to the extent observed (Figure 4f). This is an underestimate of the actual co-segregation bias because some developmental mortality likely stems from factors other than homozygosity and because we may have underestimated the true crossover rate. Thus, strongly non-Mendelian co-segregation of crossover products maintains heterozygosity in *O. biroi*.

## Discussion

Understanding extreme modes of reproduction can provide unexpected insights into fundamental cellular processes. Here, we investigated a parthenogenetic ant that produces diploid offspring by fusing two haploid products from the same meiosis (Figure 1a). This reproductive mode should lead to segmental loss of heterozygosity distal to crossovers, but we show that such loss of heterozygosity is exceedingly rare (Figure 1e, Figure 3). Surprisingly, we found that crossovers occur every meiosis, but heterozygosity is maintained because both crossover products are regularly co-inherited (Figure 2, Figure 3). This co-segregation of crossover products is a substantial departure from Mendelian segregation (Figure 4f).

The best-known form of non-Mendelian segregation is selfish meiotic drive^2^, in which driving alleles promote their own transmission over alternative alleles on the homologous chromosome. However, *O. biroi* demonstrates non-Mendelian segregation across all chromosomes without allele-specific bias. Therefore, this phenomenon likely results from a systemic perturbation of the meiotic program, where co-segregation of crossover products leads to the programmed inheritance of both alleles at all loci (Figure 4g).

As far as we know, there are only two other known cases of meiotic drive for crossover status rather than allelic identity. First, co-segregation of crossover products also occurs in the asexual nematode *Mesorhabditis belari*^24^, implying that crossover co-segregation maintains heterozygosity in distantly related lineages that have evolved parthenogenesis independently. Second, in meiosis II of human oogenesis, crossover products are nearly twice as likely as non-crossover products to end up in the oocyte rather than the second polar body^25^, a phenomenon that has been called ‘unselfish’ meiotic drive^26^. This illustrates that crossover co-segregation can also occur in sexual species, even if segregation bias is much weaker than what we report here for *O. biroi*. Given that very few studies have had the ability to detect non-random inheritance of crossovers, this raises the possibility that cellular “memory” of crossovers is a poorly understood yet widely spread aspect of “normal” chromosomal segregation. The underlying mechanisms can then be elaborated on in extreme reproductive systems such as parthenogenesis.

## Methods

### Chromosome squashes

To study chiasmata in *O. biroi* meiosis, we produced squashes of meiotic nuclei at metaphase I. For this, we slightly modified the method of Rosin et al., 2021^27^. Hymenopteran oocytes are arrested at metaphase I prior to egg laying^28,29^, so we dissected ovaries in 1x PBS, targeting the most mature oocytes available, and incubated oocytes in hypotonic solution (1% sodium citrate) on a poly-L-lysine coated glass slide for 12 minutes. Next, we removed the hypotonic solution and replaced it with 10 µl of 45% acetic acid / 1% PFA / 1x PBS for six minutes. We then physically squashed oocytes with a glass coverslip and flash-froze the slide/coverslip in liquid nitrogen. Immediately after removal from liquid nitrogen, we removed coverslips with a razor blade and incubated slides in cold 3:1 methanol:glacial acetic acid for 10 minutes. We rinsed slides twice in 1x PBS and then dehydrated them for five minutes each in 70%, 80%, and 100% ethanol (all ice-cold) before air-drying for at least 24 hours.

We stained slides with DAPI (1 µg per ml) for 10 minutes before air-drying, then added SlowFade Diamond mounting medium, added the coverslip, and sealed with nail polish. We kept mounted slides at room temperature overnight before imaging and subsequently stored them at 4°C.

We performed confocal microscopy using Zen image acquisition software on a Zeiss LSM 900 with the 405nm laser line. Images were obtained with Airyscan in super-resolution mode using a Zeiss LD LCI Plan-Apochromat 40X / 1.2NA objective lens and Zeiss Immersol Sil 406 immersion medium. We performed deconvolution using Huygens Professional version 22.04 (Scientific Volume Imaging, The Netherlands, http://svi.nl) with default parameters. Z-projection images were produced from stacks taken at 0.18µm steps using ImageJ/FIJI^30^.

### Probability of segmental loss of heterozygosity

To estimate the number of chromosomes expected to lose heterozygosity each meiosis, we used a binomial model assuming one crossover per chromosome (higher numbers of crossovers per chromosome produce a higher probability of loss of heterozygosity, meaning that, if anything, our calculations should underestimate loss of heterozygosity rates). Because all pairs of homologous chromosomes assort independently, each chromosome can be considered an independent trial (n=14 chromosomes in the *O. biroi* karyotype). Chromatids are expected to segregate randomly in meiosis II, so in central fusion automixis, there is a probability of 0.5 of crossover-induced loss of heterozygosity on any one chromosome (Figure 1b). We used the dbinom() function in R 4.0.5^31^ to calculate the binomial probability mass function for losing heterozygosity on all possible numbers of chromosomes in any given meiosis. To estimate the proportion of offspring expected to have a loss of heterozygosity, we took the sum of the probabilities that loss of heterozygosity occurs on one or more chromosomes.

To fit this model using empirical data, we substituted the empirical crossover estimate obtained from linked read sequencing (see available code). To explore the potential of co-segregation bias (i.e., the possibility that two or zero crossover products are inherited more commonly than one crossover product and one non-crossover product) to explain heterozygosity maintenance, we varied the probability of loss of heterozygosity following a crossover between 0.5 (Mendelian) and 0 (complete co-segregation bias). To explore the degree to which homozygous lethality might help maintain heterozygosity, we subtracted hypothetical developmental mortality rates ranging from 0 to 1 from the proportion of offspring expected to have a loss of heterozygosity, making the conservative assumptions that 1) all mortality comes from loss of heterozygosity, and 2) any loss of heterozygosity is lethal. To determine which parameter sets of these values are likely to reflect reality, we compared these calculations to empirically determined values for the rate of developmental mortality and the proportion of offspring with segmental losses of heterozygosity.

### Generation of the transgenic marker line

To track pedigrees of ants within colonies, we generated a transgenic line with a stable fluorescent label. For this, we used a plasmid with a piggyBac backbone to transpose the baculovirus-derived ie1 enhancer/promoter element to drive expression of the red fluorescent protein dsRed (Supplementary Figure 2). This plasmid (pBac-ie1-dsRed) was assembled as part of a previous study^32^. For transgenesis, we collected 660 eggs from clonal line B^13,33^, injected them with pBac-ie1-dsRed, and reared the resulting fluorescent individuals according to our established protocol^32^, yielding isogenic colonies of fluorescent individuals derived from single founders. The results from the injection experiments are summarized in Supplementary Table 1, and the procedure for generating and rearing transgenic ants is outlined in Supplementary Figure 2.

dsRed fluorescence in the transgenic line is detectable starting in the first larval instar and remains visible throughout metamorphosis and in adults (Supplementary Figure 2). To reduce the movement of animals during imaging, we placed them at -20°C for 5 minutes beforehand. In larvae, fluorescent dots are initially visible on each body segment. dsRed fluorescence becomes diffusely visible throughout the body during the final larval instar. This fluorescence remains visible after pupation. Pigmentation of the cuticle partially masks fluorescence in older pupae, but fluorescence is still apparent at the tip of the gaster, joints, and other areas. Fluorescence remains detectable in live adults over six months old.

To characterize the fluorescent phenotype more closely, we analyzed the fluorescence of five transgenic and five wild-type individuals, each removed from cycling colonies approximately one month after eclosion. We placed individuals in ethanol for 10 seconds before dissecting them in PBS. We dissected brains and cut bodies between the thorax and gaster. We fixed brains and gasters in 4% PFA for 30 minutes, washed them 3x in PBS with 0.1% Triton, and mounted them in Dako mounting medium. We detected dsRed and autofluorescence by imaging brains and gasters using the red and green fluorescence channels of an epifluorescence microscope (Supplementary Figure 2). While individuals from a previous study with [ie1-dsRed, ObirOrco-QF2, 15xQUAS-GcaMP6s] express dsRed in their antennal lobes^32^, fluorescence is not visible in dissected brains of ants with ie1-dsRed alone. This suggests that the ie1 promoter does not normally drive expression in brain tissue, but interaction between adjacent transgenes can lead to leaky expression^32^. The brightest dsRed fluorescence was detected in the gaster. We observed intense dsRed fluorescence in the poison gland filament, the Malpighian tubules, and structures associated with the fat body (Supplementary Figure 2). We reared a large population of a single transgenic line, which readily forms functioning colonies and produces offspring with stable transgene expression, indicating that the individuals are fully viable.

The transgenic ants developed for this study are available upon request depending on availability, and assuming that all regulatory requirements are met.

### Fitness of the transgenic marker line

To assess the possibility of a fitness deficit due to the transgenic insertion, we tested whether the transgenic ants had reduced reproductive output compared to wild-type ants of the same clonal line. We calculated the proportion of offspring with dsRed fluorescence from colonies (n=17) containing a single transgenic regular worker, and 19 wild-type ants. We took the mean proportion of transgenic offspring across two colony cycles and tested whether the value was less than the expected proportion of 0.05 (1/20 transgenic ants in each colony, with most wild-types being regular workers rather than intercastes). The mean dsRed offspring proportion was 0.11±0.54SD, not significantly less than the expected 0.05 (Supplementary Figure 3; p=0.999, one-tailed one-sample Wilcoxon test). The transgenic line, therefore, does not have reduced fitness compared to wild-type ants from the same clonal background.

### Genomic characterization of the transgenic marker line

To count and map the positions of transgene insertions within the genome, we investigated the genome of one of the samples sequenced from a known pedigree (see below for sequencing details). We aligned reads to the *O. biroi* reference genome^12^ and separately to the transgene sequence. Next, we recorded read depth at all sites using “samtools depth -aa”^34^. To identify the read depths of well-assembled genomic regions, we excluded SNPs that resulted from errors in genome assembly by obtaining heterozygous SNPs with less than 2x the genome-wide median read depth. We compared the read depth of each site in the transgene insert to read depths at an equivalent number of randomly selected trusted diploid sites. The mean read depth of the transgene insert was approximately 0.5x that of the inferred genome-wide average. Because these sequences derive from diploid *O. biroi* workers, we conclude that the transgene is present in only a single copy (Supplementary Figure 3). As expected for piggyBac, the backbone was not incorporated into the genome.

To map the genomic locus containing the transgene insertion, we identified “junction reads” that aligned to both the *O. biroi* reference genome and the transgene insert^35,36^ using the Integrative Genomics Viewer^37^ and queried alignments by each junction read name using “samtools view”^34^. We used CLUSTAL 2.1 in the R package ‘msa’^38,39^ to perform multiple sequence alignments of the junction reads from each end of the insert and to generate consensus sequences. We used BLAST^40^ to search the reference genome for the consensus sequences, which identified the same position the junction reads had aligned to. The position identified contained a ‘TTAA’ motif, which is expected at the piggyBac transposon insertion site^41–44^. The insertion was located between positions 8810280 and 8810281 on the ninth chromosomal scaffold, which is not within any predicted gene model (Supplementary Figure 3).

### Ant rearing and obtaining mother-daughter pairs

To study the amount of crossover recombination and loss of heterozygosity per meiosis, we needed to sequence genomes of ants from known pedigrees. To obtain known pedigree pairs, we created experimental colonies consisting of a single focal fluorescently labeled transgenic ant with 19 wild-type ants. After colonies laid eggs and reared them to the pupal stage, we sampled red fluorescent individuals of known pedigree (mother-daughter and sister-sister pairs). Individuals from different pedigrees were sometimes compared to sample larger numbers of meioses. In these cases, we did not know the exact pedigree, but because the ants were from a single transgenic line recently derived from a single founding individual, we knew the range of possible numbers of meioses separating each pair of individuals.

Haploid males were haphazardly collected from several stock colonies whenever they were found. To make the data as comparable as possible, we haphazardly sampled diploid females from the same stock colonies.

### Short-read whole-genome shotgun sequencing

We performed whole-genome shotgun sequencing on 22 Illumina DNA libraries. For each library, we disrupted an individual ant (adult or pupa) using a Qiagen TissueLyser II and extracted genomic DNA using Qiagen’s QIAmp DNA Micro Kit. We prepared libraries using Illumina’s Nextera DNA Flex kit and targeted 40x sequencing coverage for each library to ensure sufficient coverage to detect heterozygosity accurately. Reads were trimmed with Trimmomatic 0.36^45^, aligned with bwa mem^46^ to the *O. biroi* reference genome (Obir_v5.4, GenBank assembly accession: GCA_003672135.1), and subsequently sorted, deduplicated, and indexed with picard^47^. Variants were called with GATK HaplotypeCaller (version 4.2)^48^ and filtered using GATK’s hard-filtering recommendations.

We performed additional filtering to include only high-confidence variants in this analysis. We excluded falsely collapsed regions in the reference genome by filtering against sites found to be heterozygous in haploid males, variants with three or more alleles in a single diploid clonal lineage, and sites with read depth greater than twice the genome-wide mean. To exclude erroneous calls of heterozygosity or homozygosity, we filtered against sites with read depth of less than 15, with proportionate minor allelic depth of less than 0.25 in putatively heterozygous samples, or with nonzero minor allelic depth in putatively homozygous samples. While performing data analysis, we observed that forgoing any of these additional filtering steps led to many false positive gains of heterozygosity, losses of heterozygosity, and recombination events.

### Loss of heterozygosity in known pedigrees

To analyze mother-daughter pairs, we classified all differences in genotype as loss of heterozygosity or gain of heterozygosity. Point-mutation-induced gains of heterozygosity were inferred at sites where an individual was heterozygous and possessed an allele found in no other members of that clonal lineage. To screen against remaining false positives, we visually inspected all putative changes in heterozygosity by loading aligned reads in IGV2.6.2^37^ to remove all sites with ambiguous heterozygosity calls (i.e., putatively homozygous sites with one or more aligned reads bearing the alternate allele), or with signatures of assembly errors (variants with reads indicative of heterozygosity in haploid males). Viewing alignment files in this way included reads GATK had screened before genotype calling, and such reads often contradicted genotype calls at sites with putative changes in heterozygosity. For a global assessment of heterozygosity differences, we constructed non-overlapping 10kb windows (21784 in total) and, for each sample, recorded the proportion of such windows that contained at least one heterozygous SNP.

### Comparison of recombination among haploids and loss of heterozygosity among diploids

To analyze recombination among samples of unknown pedigree, we first inferred ancestrally heterozygous sites following Oxley et al., 2014. For each clonal line, we considered sites heterozygous in any diploid to be ancestrally heterozygous as long as 1) all other diploids had identical heterozygous genotypes or had genotypes homozygous for one of the alleles found in the heterozygous genotype, and 2) all haploids were hemizygous for one of the two alleles. Homozygous genotypes at ancestrally heterozygous sites were classified as resulting from loss of heterozygosity. Contiguous runs of such SNPs greater than 1500bp in length were classified as losses of heterozygosity resulting from crossovers, whereas shorter stretches were classified as gene conversion tracts. Changing this threshold to 5000bp (the approximate upper bound of non-crossover maximum tract length in *Drosophila melanogaster*^19^) did not qualitatively affect the results (i.e., the ratio of crossovers detected among haploids to crossovers inferred among diploids changed from 43.3 to 42.3 for Line A, and from 139.3 to 136.3 for Line B). We directly observed recombination events among the genomes of haploid males by recording changes in relative phasing among samples at ancestrally heterozygous sites.

The lengths of haplotype blocks (contiguous tracts of SNPs with identical relative phasing) were recorded as the number of bps between the first SNP and last SNP in each haplotype block. To ensure that gene conversion events did not inflate the number of crossovers detected, we removed gene conversion-sized events from the dataset before counting crossover recombination events. For the small number of losses of heterozygosity among diploid females that resulted from crossovers, we counted crossover events only at sites for which the transition from heterozygosity to homozygosity was visible, in accordance with how crossovers were inferred among male haplotypes.

To examine the correlation of crossover recombination with genetic differentiation across colonies, we calculated genetic distance via identity by state at ancestrally heterozygous SNPs using the R package SNPRelate^49^. To ensure statistical independence of comparisons, we did not perform all possible pairwise comparisons but instead compared all samples to a single reference individual. From each reference colony (i.e., Line A: C16 and Line B: STC6), we arbitrarily selected a single diploid female and haploid male as reference individuals for comparison. We counted the number of recombination events between each individual and the reference male and calculated the genetic distance between each individual and the reference diploid female. Using diploid genotypes to calculate genetic distance ensured statistical independence from the recombination events evident among haploid genomes. Because the ability to detect crossovers that have occurred over many generations scales positively with marker density, and the clonal lines differed in the number of ancestrally heterozygous SNPs, we normalized the crossover counts for each pair by dividing the number of crossovers detected by the number of ancestrally heterozygous SNPs in that clonal line.

### Linked-read sequencing

To determine whether crossovers occur every meiosis, we performed linked-read whole genome shotgun sequencing on 17 libraries. To extract high molecular weight DNA, we used Qiagen’s MagAttract kit after grinding individual ants (adult or pupal stage) under liquid nitrogen using an up-and-down motion with a plastic pestle. We prepared TELL-Seq^50^ libraries according to the manufacturer’s instructions, using two ng of input DNA, 15 µl of TELL beads, and ten amplification cycles. We targeted at least 60x genome-wide sequencing coverage for each library to ensure reliable phasing.

We performed initial data analysis using pipelines available from Universal Sequencing Technologies. We aligned and ‘linked’ reads using Tell-Read and performed variant calling and phasing using Tell-Sort^50^. This pipeline uses bwa to align reads to the reference genome, calls variants using GATK HaplotypeCaller, and phases reads using HapCUT2^51^. We removed all sites with genotype quality (GQ) less than 99 or phasing quality (PQ) less than 100. We next excluded falsely collapsed regions in the reference genome by filtering against sites found to be heterozygous in haploid males, variants with three or more alleles in a single clonal lineage, and sites with read depth less than twenty or greater than twice the genome-wide mean.

### Analysis of recombination among linked-read libraries

Putative recombination events were identified between phased vcf files using “vcftools – diff-switch-error,”^52^ which produces a list of all pairs of consecutive SNP positions between which the phasing differs. By comparing phased diploid genomes, we can only confidently infer recombination events in genomic regions reliably phased in both focal samples. Therefore, we screened out putative recombination events that could not be unambiguously confirmed. We removed putative events in which the two focal SNPs were intervened by, or were flanked by, signatures of poor genome assembly. These included sites with no aligned reads in one or more samples, poor mapping quality (aligned reads with MAPQ=0), or signatures of falsely concatenated genomic regions (coverage greater than twice the genome-wide average, or variants observed to be heterozygous in haploid males). To rule out erroneously phased genomic regions, we required that the haplotypic phase remain consistent for at least 2 SNPs upstream and downstream of putative recombination events. We visually inspected all putative recombination events in IGV.

To determine the proportion of the genome for which crossovers were detectable, we identified all possible pairs of contiguous SNPs and determined whether that pair would have been eliminated by one or more of the above filtering criteria in each pair of samples. Then, we took the proportion of base pairs that fell between retained pairs of SNPs as the proportion of the genome for which crossovers were detectable. For the 14 pairs of samples, this proportion fell between 0.50 and 0.61, indicating that the observed number of crossovers from this analysis is an underestimate by at least 39%.

### Developmental mortality estimates

One mechanism to maintain heterozygosity would be the mortality of individuals with losses of heterozygosity. Such mortality would have to occur after eggs are laid (the meiotic divisions and subsequent central fusion occur following egg-laying in *O. biroi*). Measuring developmental mortality requires tracking the survival of known numbers of eggs in replicate colonies, which is complicated by the fact that *O. biroi* are all reproductively totipotent, and thus, the caregiver ants could lay additional eggs that would lead to underestimates of developmental mortality. To avoid this complication, we created seven replicate colonies in Petri dish nests with moistened plaster of Paris floors, each with 46 fluorescently labeled transgenic eggs, 40 non-transgenic adults, and 38 non-transgenic pupae (pupae were included so that developing larvae could drink their secretions^53^; these pupae eclosed as adults during the experiment). We allowed the adults to rear the eggs, and once the eggs had hatched as larvae, we fed the colonies daily with three to five frozen fire ant pupae and cleaned and watered the plaster as needed. Once the larvae had pupated, we counted the fluorescently labeled pupae. We measured mortality at the pupal stage because all genome sequencing to determine rates of loss of heterozygosity was performed using pupal offspring.

### Data visualization

Karyoplots were drawn using karyoploteR^54^, binomial probability mass functions were drawn using base R graphics^31^, and other plots were drawn using ggplot2^55^. Illustrations and final figures were made using Adobe Illustrator.

### Statistics

We constructed linear models using the lm() function in R 4.0.5^31^. These were used to determine the slope and y-intercept of the relationship between the normalized number of crossovers and phylogenetic distance among haploid males. We also used linear models as an internal control to confirm the validity of detecting crossovers in linked read sequencing data by assessing whether the number of crossovers scaled positively with the number of meioses.

Zero of the 116 crossovers observed in our linked-read sequencing experiment were associated with losses of heterozygosity. To determine whether these data differed significantly from the Mendelian expectation that 2/3 of inherited crossovers should be associated with losses of heterozygosity, we conducted a binomial test in GraphPad Prism.

To calculate the 95% confidence interval of the proportion of offspring that bear losses of heterozygosity, we used the Wilson/Brown^56^ method in GraphPad Prism. This method is recommended for proportions very close to zero or one. We used standard methods to calculate all other 95% confidence intervals.

A one-tailed, one-sample Wilcoxon test was used to determine if the fitness of the transgenic line was less than that of wild-type ants.

## Data availability

All DNA sequencing data are publicly available at the National Center for Biotechnology Information Sequence Read Archive under accession number PRJNA947942.

## Code availability

All code is available on GitHub (https://github.com/kipdlacy/UnselfishMeioticDrive).

## Acknowledgments

We thank Stephany Valdés-Rodríguez and Leonora Olivos-Cisneros for help obtaining ants for sequencing; Ivan Lacroix for help with ovary dissections; the entire Kronauer Laboratory for helpful discussion and for contributing male ants; Connie Zhao and the Rockefeller University Genomics Resource Center for genome sequencing and helpful advice; Jen Balacco and Olivier Fedrigo at the Rockefeller University Reference Genome Resource Center for helpful discussion and use of the Femto Pulse; Priyam Banerjee at the Rockefeller University Bioimaging Resource Center for advice and training; Buck Trible for the suggestion to inspect genomics data visually in IGV; Orli Snir, Hironori Funabiki, Li Zhao, Fred Cross, Agata Smogorzewska, participants of the 2022 Meiosis GRC, as well as Tom Carroll and the Rockefeller Bioinformatics Resource Center for helpful discussion; Molly Nelson for help drawing ant cartoons. This work was supported by a Gabrielle H. Reem and Herbert J. Kayden Early-Career Innovation Award and the National Institute of General Medical Sciences of the National Institutes of Health under Award Number R35GM127007, both to D.J.C.K. The content is solely the responsibility of the authors and does not necessarily represent the official views of the National Institutes of Health. D.J.C.K. is an Investigator of the Howard Hughes Medical Institute. This is Clonal Raider Ant Project paper #29.

## Contributions

K.D.L. and D.J.C.K. designed research; T.H. created the transgenic line, and T.H. and K.D.L. characterized the line; K.D.L. performed research, analyzed data, and visualized results; K.D.L. and D.J.C.K. wrote the manuscript; D.J.C.K. supervised the project.

## Supplementary Figures & Tables

**Supplementary Figure 1.**
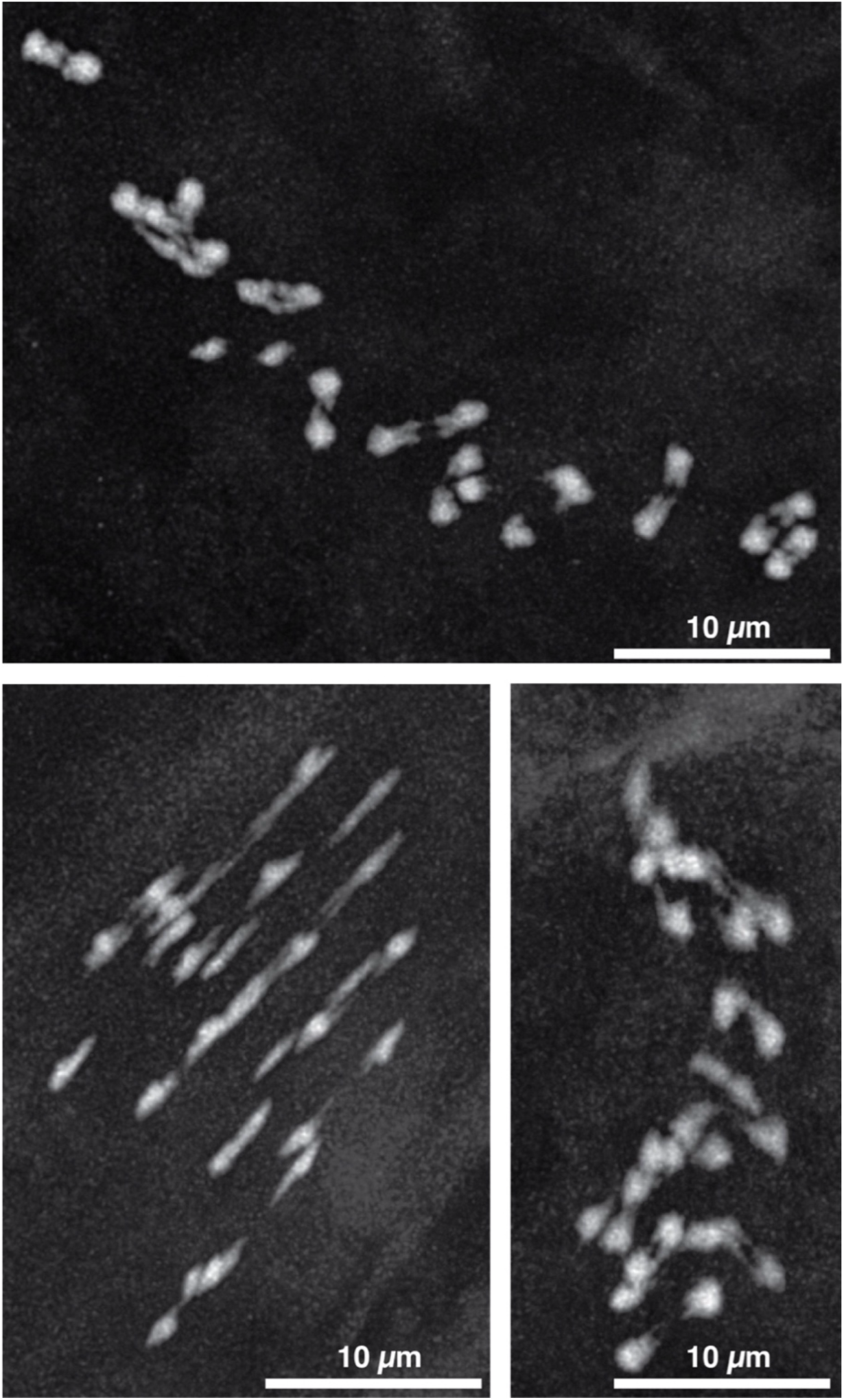
Additional metaphase I spreads reveal that chiasmata connect paired homologs. Maximum projections of DAPI-stained *O. biroi* chromosomes at metaphase I reveal DAPI connections between paired homologs that indicate chiasmata. Each projection is from a different oocyte produced by a different ant.

**Supplementary Figure 2.**
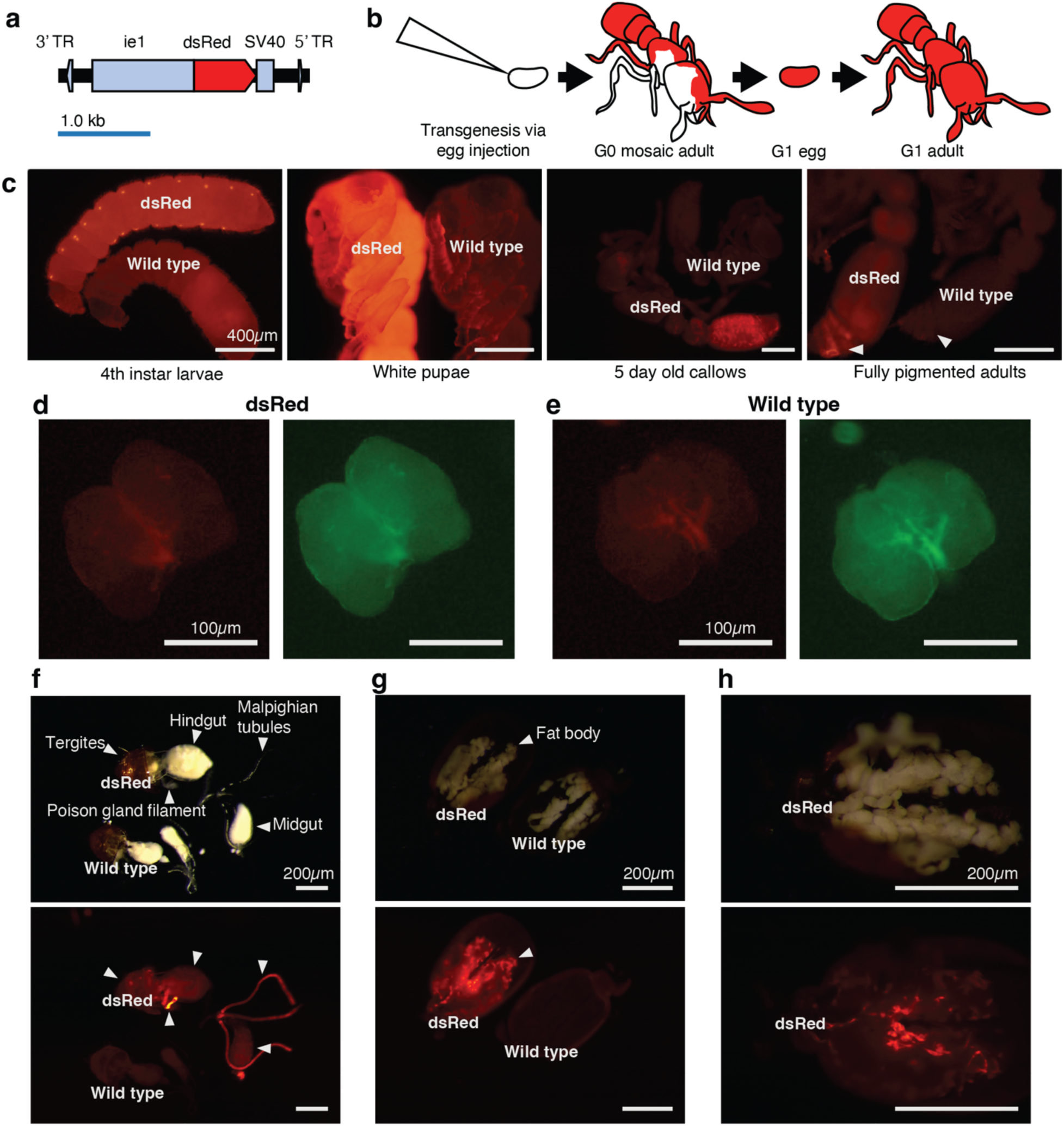
Generating the transgenic line. **a,** Diagram of the transgene construct pBac-ie1-dsRed. Promoter/enhancer ie1, dsRed coding sequence, and SV40 termination sequence were assembled between the 3’ and 5’ Terminal Repeats (TRs) on a piggyBac plasmid backbone. **b,** Schematic for generating transgenic clonal raider ants. Young eggs are collected from wild-type (WT) ants and injected with transposase/vector mix. Some eggs develop into G0s with somatic mosaicism of the integrated transposon. G0s are reared in groups and screened for fluorescence. Eggs are collected from the mixed population of G0s and reared to produce G1s with potentially unique transgene integrations. **c,** Fluorescent phenotypes of transgenic clonal raider ants. In each panel, a transgenic individual from clonal line B is shown next to a WT individual from clonal line B at the same developmental stage, imaged under epifluorescence. Left to right: fourth instar larvae, pupae, five-day-old callows, and fully pigmented adults. In the rightmost panel, the arrows indicate the region of brightest fluorescence in dsRed individuals and the corresponding spot in WTs, respectively. Fluorescence in transgenic individuals is visible in live individuals across all life stages, excluding eggs. **d,e,** Brains from transgenic (**d**) and WT clonal line B (**e**) ants were dissected and imaged under epifluorescence. Only autofluorescence is visible in brains, revealing that dsRed is not expressed in brains in this transgenic line. Autofluorescence in the green channel is shown for each image to visualize anatomy (right panels in **d,e,**). **f,** One transgenic gaster and one WT clonal line B gaster were dissected side by side, both from 5-day-old callows. Dissected tissues are visible in bright field (top), and dsRed fluorescence is visible under epifluorescence in several tissues, including the poison gland filament and Malpighian tubules (bottom). **g,** Fat body tissue attached to the cuticle of the gaster is visible under bright field in transgenic and WT individuals (top), with dsRed detected in structures associated with the fat body (bottom). **h,** Close-up on the transgenic fat body from (g).

**Supplementary Figure 3.**
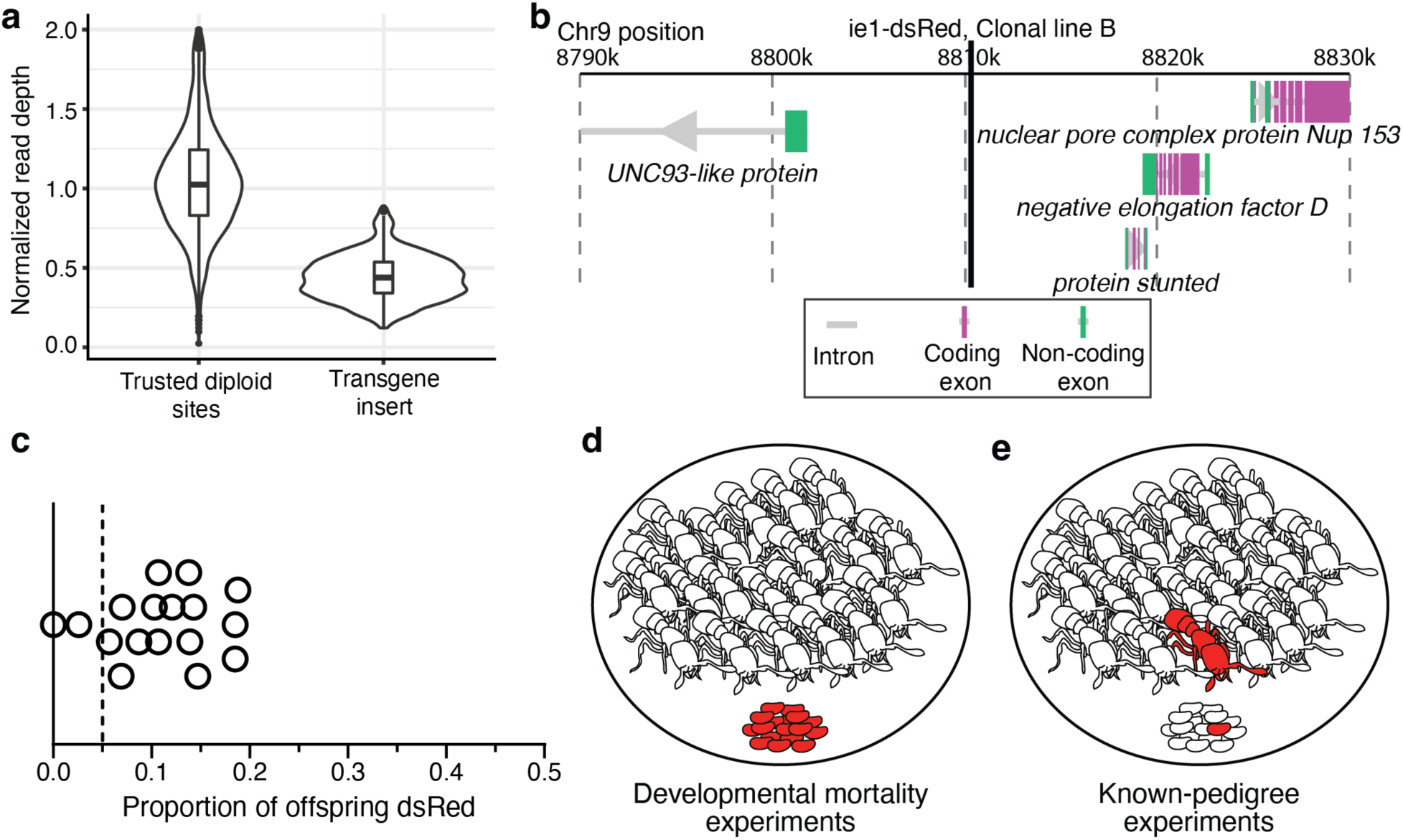
Characterizing and utilizing the transgenic line. **a,** Normalized read depth at a panel of 2,628 trusted diploid heterozygous sites in the *O. biroi* reference genome compared to read depth at all sites in the 2,628bp transgene insert sequence based on whole genome resequencing of a single transgenic ant. The read depth at the transgene insert was approximately 0.5x the read depth at heterozygous SNPs in the *O. biroi* reference genome, revealing that a single insertion occurred. Box plots show median (center line), interquartile range (IQR) (box limits), and 1.5 x IQR (whiskers). Data points that fall outside 1.5 x IQR are shown individually. Violin plots show the kernel probability density, meaning that the proportion of the data located at a y-axis value is represented by the width of the outlined area. **b,** Genomic context of the mapped insertion site for the transgene on the ninth chromosomal scaffold of the *O. biroi* reference genome. The insertion did not interrupt any annotated genes. **c,** The transgenic line does not have a fitness deficit. Replicate colonies with one transgenic individual and 19 wildtype individuals were allowed to cycle, and the proportion of transgenic offspring were counted. The vertical dotted line indicates the expected proportion of transgenic offspring. **d,e,** Cartoons of the experimental setups for developmental mortality experiments **d** and known-pedigree-producing colonies **e**. **d,** For developmental mortality experiments, replicate colonies contained 46 fluorescently labeled eggs, 40 WT adults, and 38 WT pupae (not shown). By counting the number of fluorescent offspring after eggs had developed, we determined mortality rates while avoiding the possibility of underestimation due to additional egg-laying by adults throughout the experiment. **e,** For known-pedigree production, we housed one fluorescently labeled ant with 19 WT ants and allowed colonies to lay eggs and rear their offspring. We sequenced whole genomes of fluorescently labeled pairs of mothers and daughters or pairs of sisters. This experimental setup was also used for the fitness experiments shown in **c**.

**Supplementary Figure 4.**
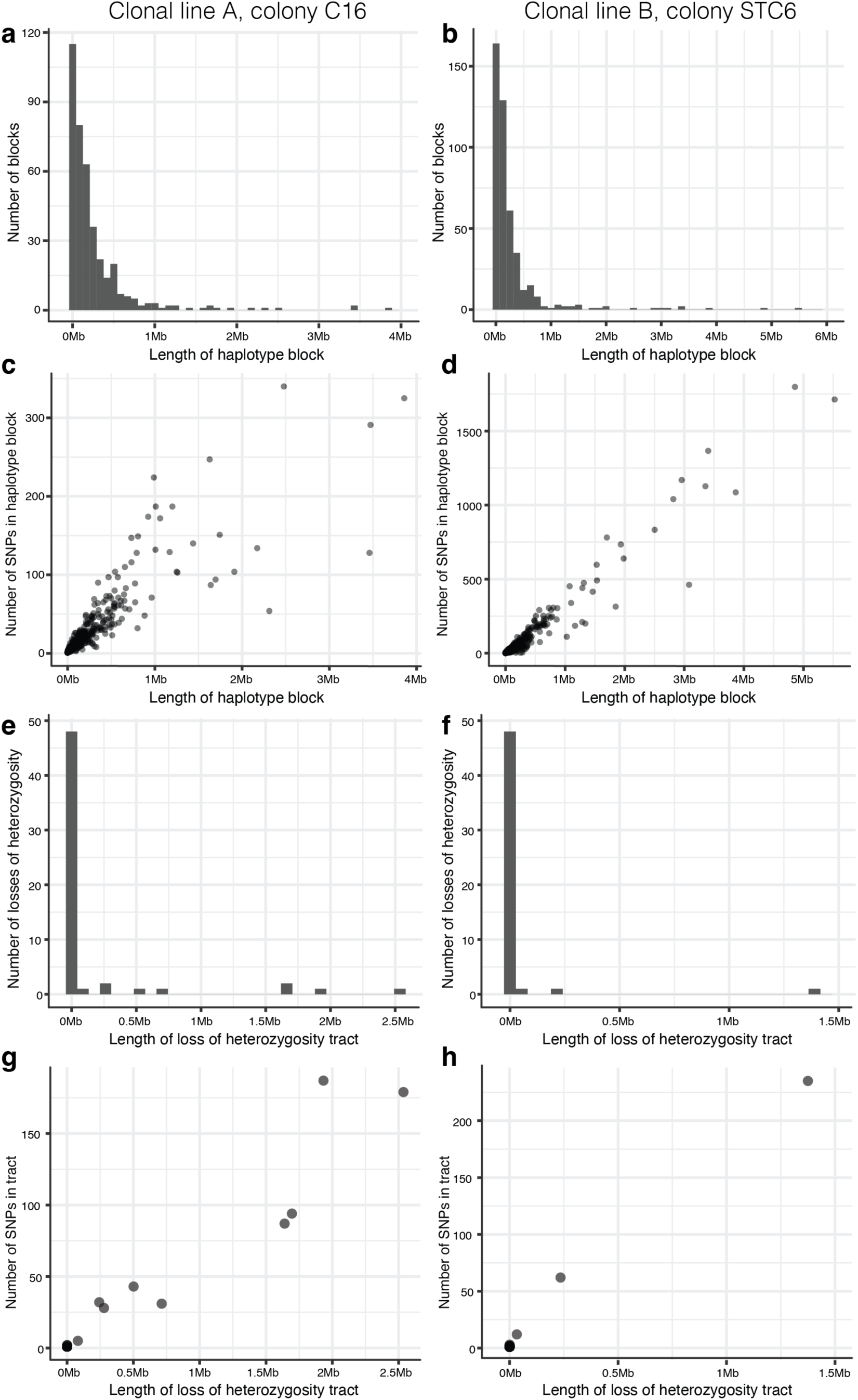
Haplotype blocks and loss of heterozygosity tracts resulting from crossovers and gene conversion within colonies of *O. biroi*. **a,b,c,d,** The distribution of haplotype block lengths among haploid males and **e,f,g,h,** the distribution of sizes of loss of heterozygosity among diploid females from a single colony of clonal line A (**a,c,e,g**; colony C16) and clonal line B (**b,d,f,h,**; colony STC6). **a,b,** Histograms depict the distance in base pairs between recombination events, or between a recombination event and the end of the contig on which it occurred. In addition to gene conversion events, which span at most a few kb in length, short haplotypes represent remnants of comparatively old recombination events that many subsequent recombination events have since broken up. By contrast, long haplotype blocks likely were formed by relatively recent recombination events and have yet to be broken up by additional recombination events. The fact that many intermediate sizes of haplotypes are found in this dataset suggests that the observed recombination events occurred gradually over many generations. **c,d,** Scatterplots depicting the association between the length of haplotype blocks and the number of ancestrally heterozygous SNPs comprising each haplotype block. Each point represents one of the haplotype blocks also depicted in **a** and **b**. Long haplotype blocks inferred to have resulted from crossover recombination were supported by a corresponding number of SNPs. **e,f,** Histograms depict the length of loss of heterozygosity tracts or distances between the beginning of a loss of heterozygosity tract and the end of a contig. Note that the total number of tracts of losses of heterozygosity is greater than the number of crossovers reported in the main text because long segmental losses of heterozygosity that spanned multiple contigs likely resulted from a smaller number of crossover events (see Methods). **g,h,** Scatterplots depict the number of ancestrally heterozygous SNPs that became homozygous as a function of the length of the respective loss of heterozygosity tract. Each point represents one of the haplotype blocks also depicted in **e** and **f**.

**Supplementary Figure 5.**
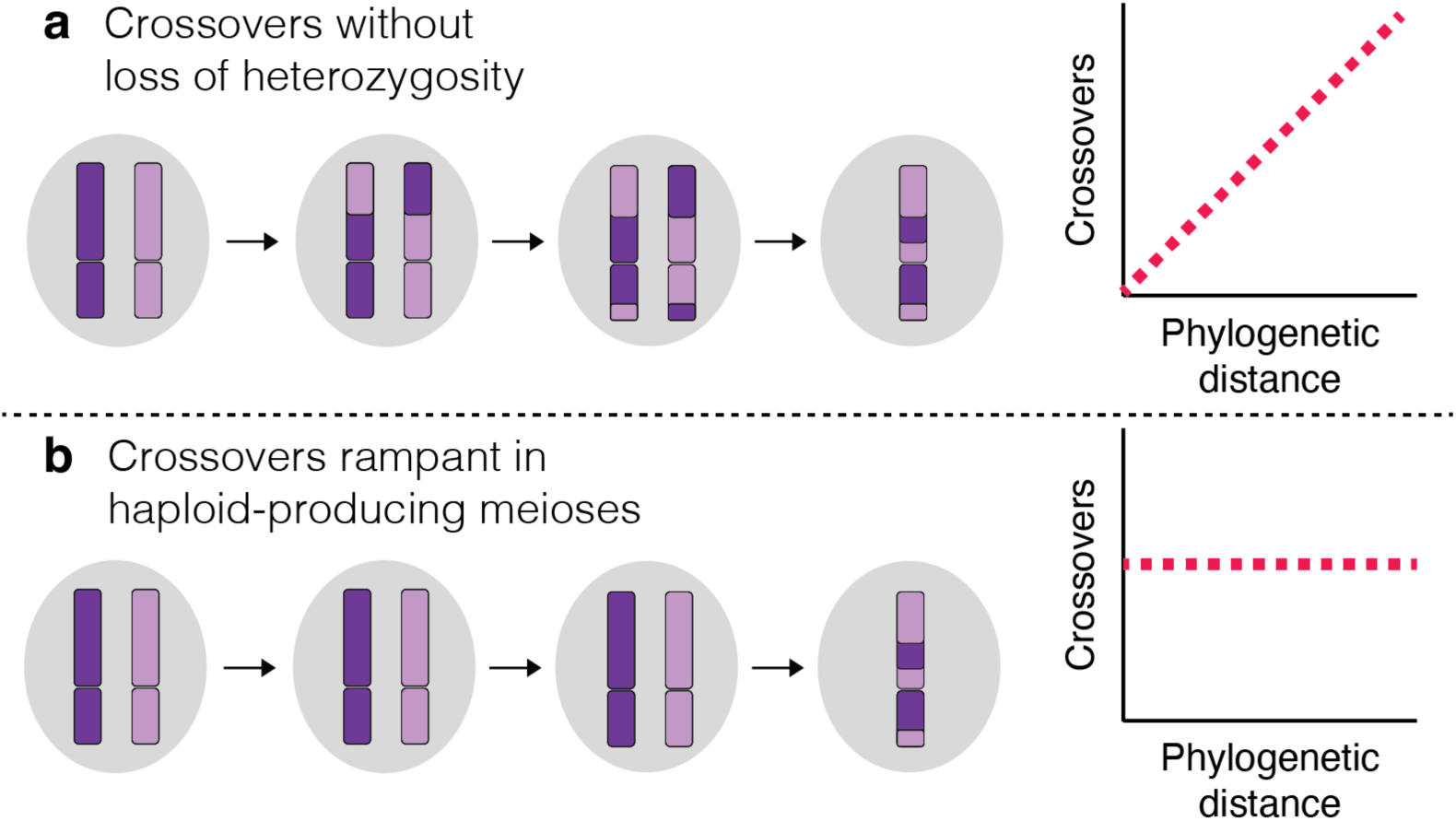
Alternative explanations for the elevated number of crossovers detectable among haploid genomes relative to diploid genomes. Illustrations of alternative explanations for the discrepancy in the number of crossovers detectable in haploid versus diploid genomes and predictions of how the number of crossovers should scale with phylogenetic distance for each explanation. Purple coloring on chromosomes indicates allelic identity. Red dotted lines in plots indicate hypothesized trends for each scenario. **a,** If crossovers occur in typical *O. biroi* meioses that are followed by central fusion and produce diploid offspring, then crossovers accumulate slowly over generations. If true, crossovers would also be present in diploid genomes but are undetectable because heterozygosity is not lost. In this scenario, the number of crossovers detected between pairs of haploids would scale positively with the phylogenetic distance between those individuals and would have a y-intercept close to zero. **b,** If crossovers were absent from typical *O. biroi* meioses, but crossovers were rampant in meioses that are not followed by central fusion (the “haploid-producing” meioses), then similar numbers of crossovers should be detectable among pairs of haploid males, regardless of genetic distance. In this scenario, the number of crossovers detected between pairs of haploids would have a slope of zero and a positive y-intercept.

**Supplementary Figure 6.**
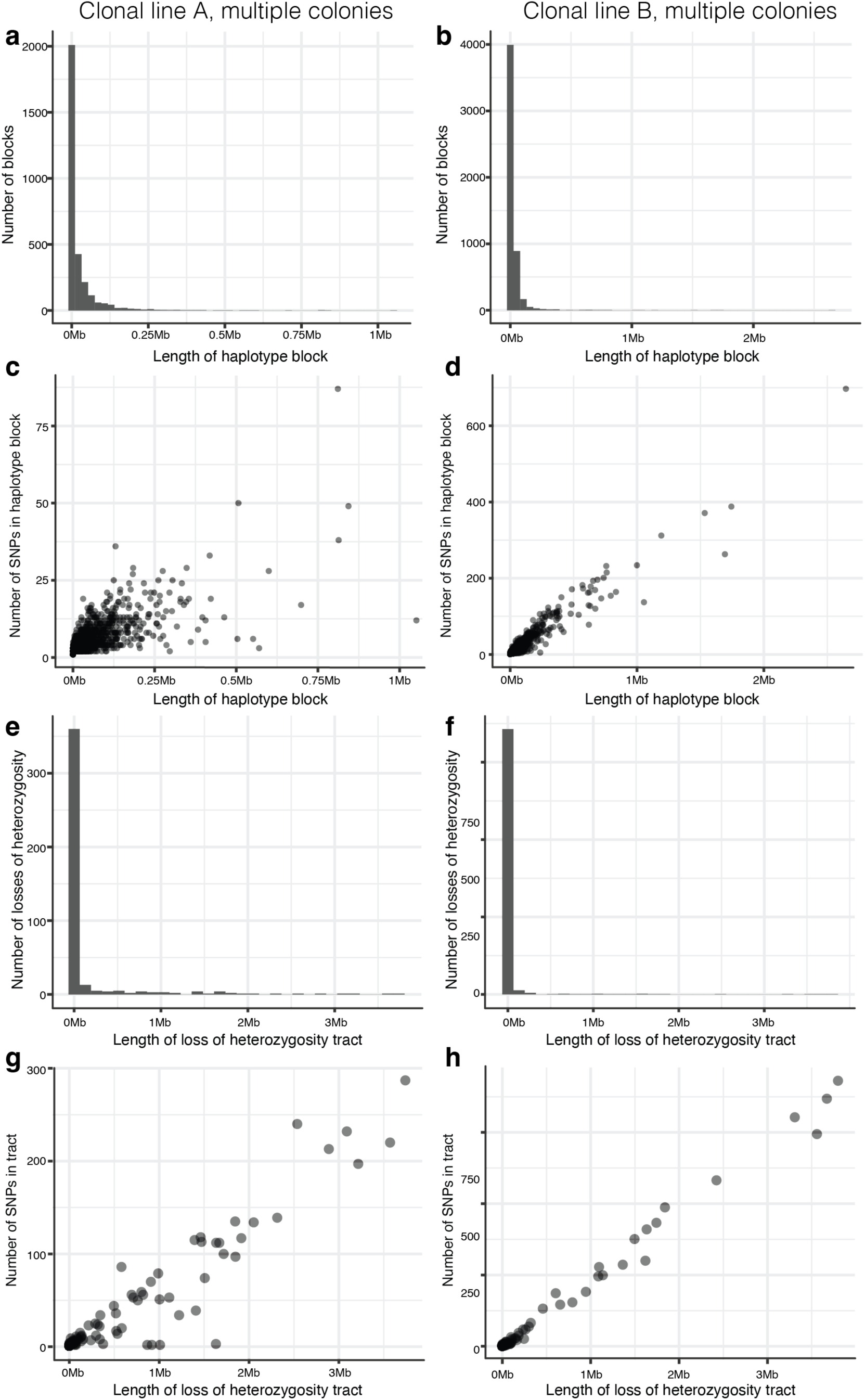
Haplotype blocks and loss of heterozygosity tracts resulting from crossovers and gene conversion across different colonies of *O. biroi*. **a,b,c,d,** The distribution of haplotype block lengths among haploid males and **e,f,g,h,** of size of loss of heterozygosity among diploid females from multi-colony datasets of Line A **(a,c,e,g)** and Line B **(b,d,f,h)**. **a,b,** Histograms depict the distance in base pairs between recombination events, or between a recombination event and the end of the contig on which it occurred. In addition to gene conversion events, which span at most a few kb in length, short haplotypes represent remnants of comparatively old recombination events, which many subsequent recombination events have since broken up. By contrast, long haplotype blocks likely were formed by relatively recent recombination events and have yet to be broken up by additional recombination events. **c,d,** Scatterplots depicting the association between the length of haplotype blocks and the number of ancestrally heterozygous SNPs comprising each haplotype block. Each point represents one of the haplotype blocks also depicted in **a** and **b**. **e,f,** Histograms depict the length of loss of heterozygosity tracts, or distances between the beginning of a loss of heterozygosity tract and the end of a contig. Note that the total number of tracts of losses of heterozygosity is greater than the number of crossovers reported in the main text because long losses of heterozygosity that spanned multiple contigs likely resulted from a smaller number of crossover events (see Methods). **g,h,** Scatterplots depict the number of SNPs that were ancestrally heterozygous but became homozygous as a function of the length of the respective loss of heterozygosity tract. Each point represents one of the haplotype blocks also depicted in **e** and **f**.

**Supplementary Figure 7.**
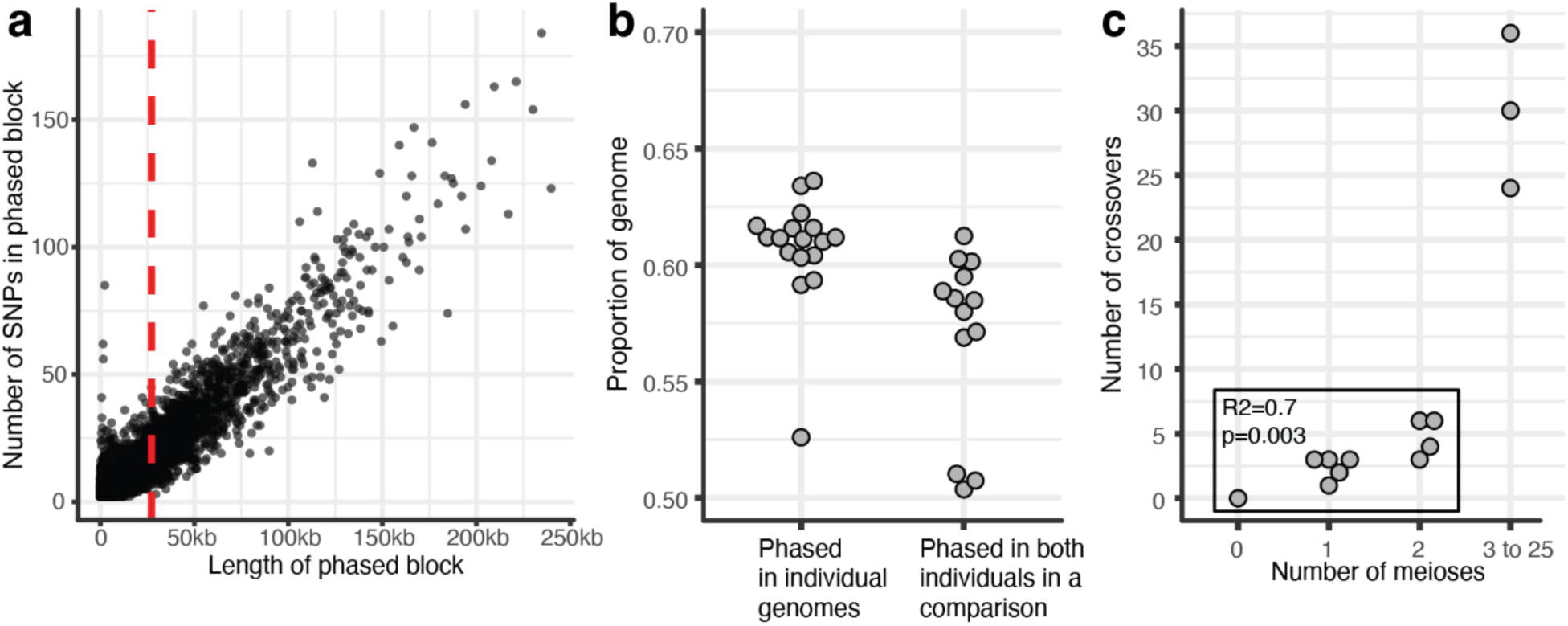
Phasing linked-read sequencing libraries to detect crossovers. **a,** The relative phasing efficiency for a single example individual library (LinkedReadPedigree3_Daughter_1) is given by a scatterplot depicting the association between the length of phased blocks and the number of ancestrally heterozygous SNPs per block, with the red line representing the phased block N50. **b,** The proportion of the genome that was successfully phased is given for all individual linked-read genomes and for all pairs of individuals compared for crossover detection. **c,** Scatterplot depicting the association between the number of crossovers detected and the number of intervening meioses for each pair of genomes. Zero meioses separate two libraries prepared from the same individual (one negative control); one meiosis separates mothers and daughters in five pairs, and two meioses separate sisters in four pairs. The pedigree was not known for three “positive control” comparisons, but the number of intervening meioses falls between 3 and 25 (see Methods). The correlation between the number of crossovers and the number of meioses is given for all known-pedigree comparisons (box).

**Supplementary Table 1.**
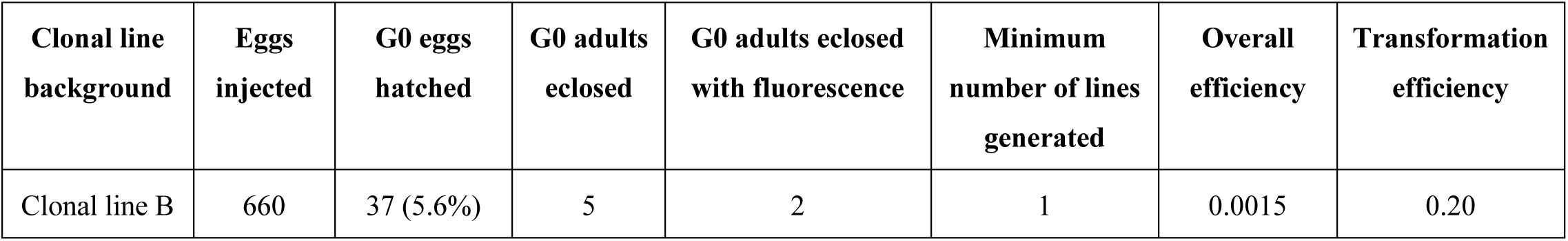
Transgenesis efficiency. Overall efficiency was calculated as the number of independent lines generated divided by the number of embryos injected. Transformation efficiency was calculated as the number of independent lines divided by the number of eclosed G0s.

| Clonal line background | Eggs injected | G0 eggs hatched | G0 adults eclosed | G0 adults eclosed with fluorescence | Minimum number of lines generated | Overall efficiency | Transformation efficiency |
| --- | --- | --- | --- | --- | --- | --- | --- |
| Clonal line B | 660 | 37 (5.6%) | 5 | 2 | 1 | 0.0015 | 0.20 |

**Supplementary Table 2.**
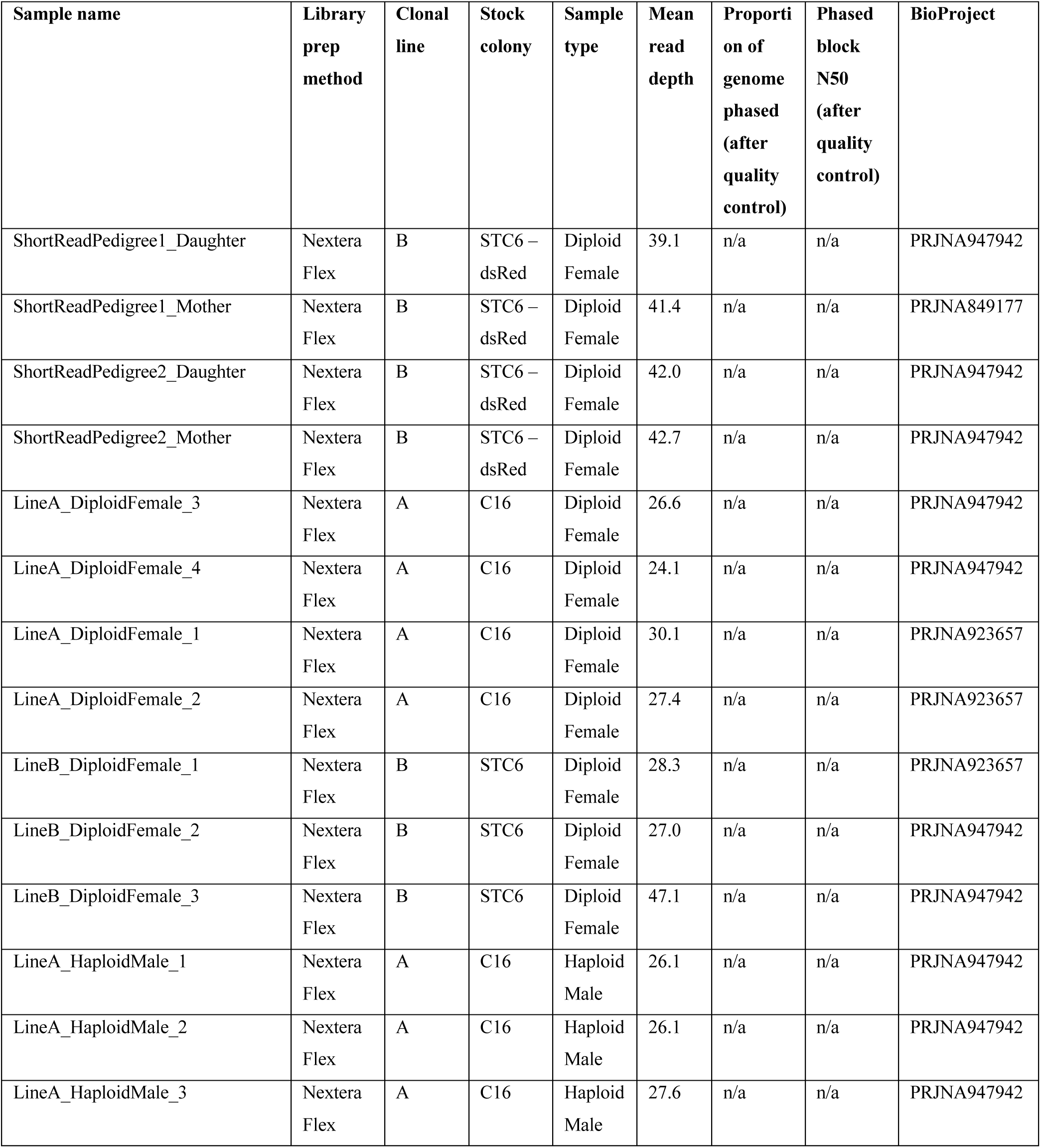
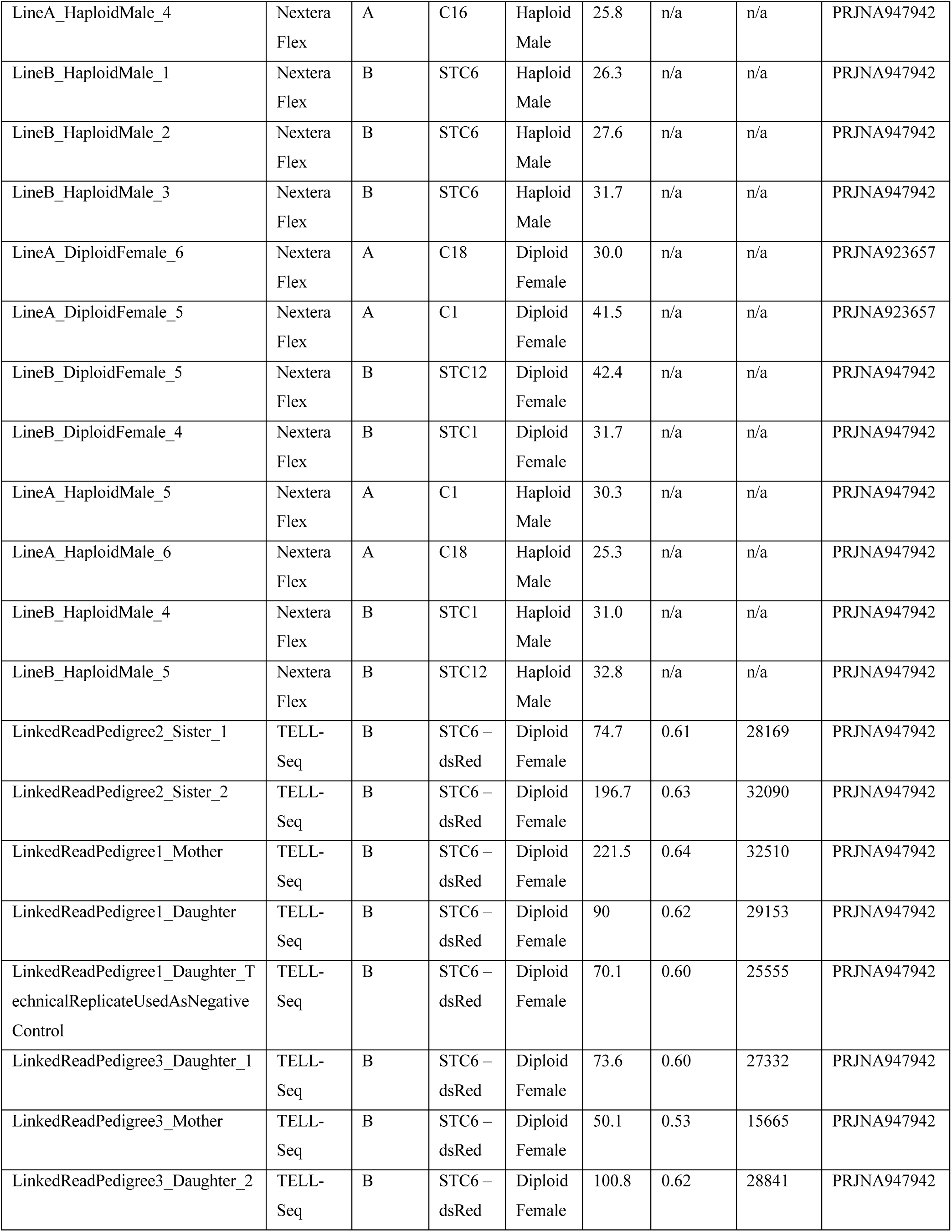
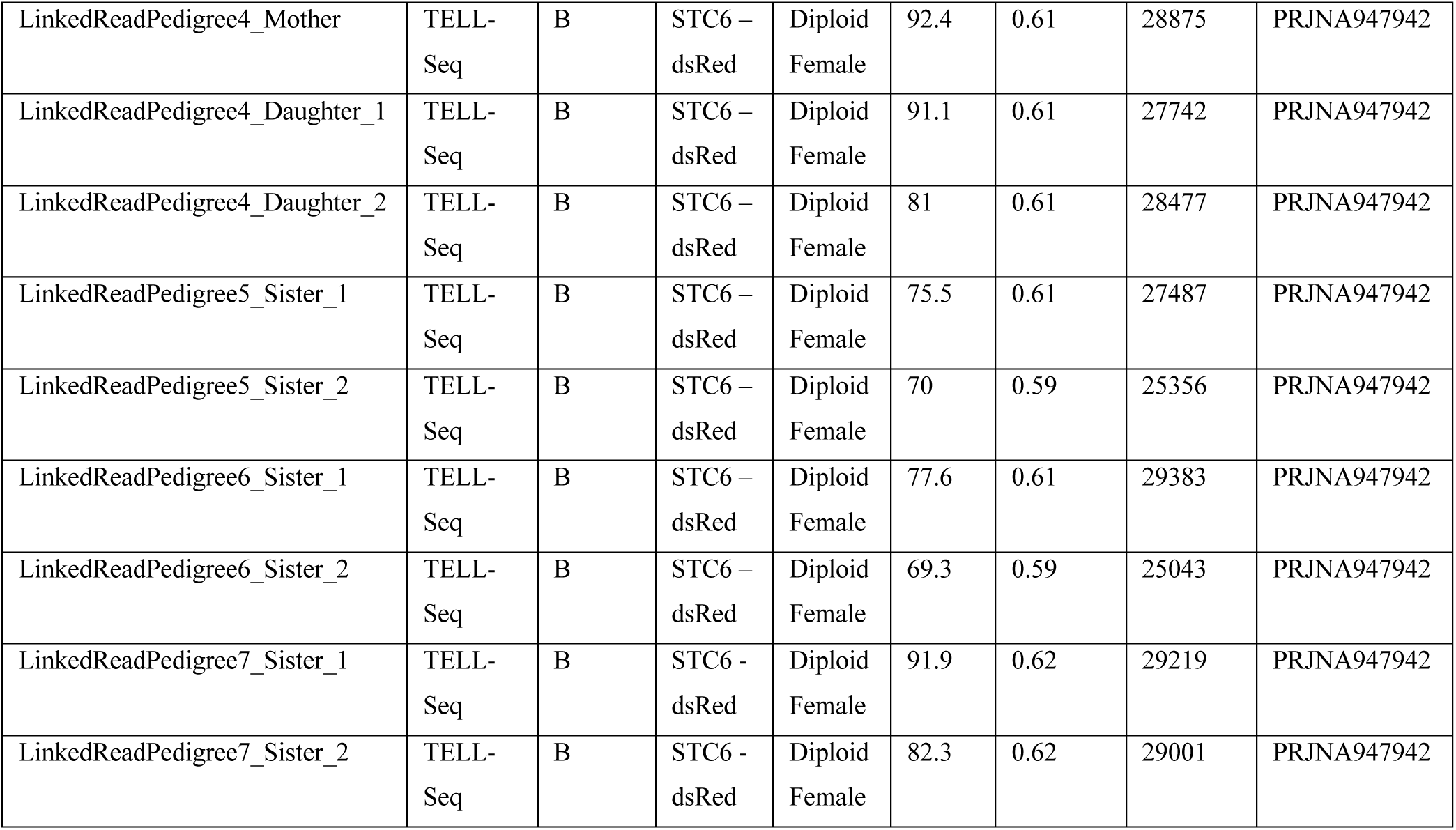
Metadata for all DNA whole-genome shotgun sequencing libraries included in this study.

| Sample name | Library prep method | Clonal line | Stock colony | Sample type | Mean read depth | Proportion of genome phased (after quality control) | Phased block N50 (after quality control) | BioProject |
| --- | --- | --- | --- | --- | --- | --- | --- | --- |
| ShortReadPedigree1_Daughter | Nextera Flex | B | STC6 – dsRed | Diploid Female | 39.1 | n/a | n/a | PRJNA947942 |
| ShortReadPedigree1_Mother | Nextera Flex | B | STC6 – dsRed | Diploid Female | 41.4 | n/a | n/a | PRJNA849177 |
| ShortReadPedigree2_Daughter | Nextera Flex | B | STC6 – dsRed | Diploid Female | 42.0 | n/a | n/a | PRJNA947942 |
| ShortReadPedigree2_Mother | Nextera Flex | B | STC6 – dsRed | Diploid Female | 42.7 | n/a | n/a | PRJNA947942 |
| LineA_DiploidFemale_3 | Nextera Flex | A | C16 | Diploid Female | 26.6 | n/a | n/a | PRJNA947942 |
| LineA_DiploidFemale_4 | Nextera Flex | A | C16 | Diploid Female | 24.1 | n/a | n/a | PRJNA947942 |
| LineA_DiploidFemale_1 | Nextera Flex | A | C16 | Diploid Female | 30.1 | n/a | n/a | PRJNA923657 |
| LineA_DiploidFemale_2 | Nextera Flex | A | C16 | Diploid Female | 27.4 | n/a | n/a | PRJNA923657 |
| LineB_DiploidFemale_1 | Nextera Flex | B | STC6 | Diploid Female | 28.3 | n/a | n/a | PRJNA923657 |
| LineB_DiploidFemale_2 | Nextera Flex | B | STC6 | Diploid Female | 27.0 | n/a | n/a | PRJNA947942 |
| LineB_DiploidFemale_3 | Nextera Flex | B | STC6 | Diploid Female | 47.1 | n/a | n/a | PRJNA947942 |
| LineA_HaploidMale_1 | Nextera Flex | A | C16 | Haploid Male | 26.1 | n/a | n/a | PRJNA947942 |
| LineA_HaploidMale_2 | Nextera Flex | A | C16 | Haploid Male | 26.1 | n/a | n/a | PRJNA947942 |
| LineA_HaploidMale_3 | Nextera Flex | A | C16 | Haploid Male | 27.6 | n/a | n/a | PRJNA947942 |
| LineA_HaploidMale_4 | Nextera Flex | A | C16 | Haploid Male | 25.8 | n/a | n/a | PRJNA947942 |
| LineB_HaploidMale_1 | Nextera Flex | B | STC6 | Haploid Male | 26.3 | n/a | n/a | PRJNA947942 |
| LineB_HaploidMale_2 | Nextera Flex | B | STC6 | Haploid Male | 27.6 | n/a | n/a | PRJNA947942 |
| LineB_HaploidMale_3 | Nextera Flex | B | STC6 | Haploid Male | 31.7 | n/a | n/a | PRJNA947942 |
| LineA_DiploidFemale_6 | Nextera Flex | A | C18 | Diploid Female | 30.0 | n/a | n/a | PRJNA923657 |
| LineA_DiploidFemale_5 | Nextera Flex | A | C1 | Diploid Female | 41.5 | n/a | n/a | PRJNA923657 |
| LineB_DiploidFemale_5 | Nextera Flex | B | STC12 | Diploid Female | 42.4 | n/a | n/a | PRJNA947942 |
| LineB_DiploidFemale_4 | Nextera Flex | B | STC1 | Diploid Female | 31.7 | n/a | n/a | PRJNA947942 |
| LineA_HaploidMale_5 | Nextera Flex | A | C1 | Haploid Male | 30.3 | n/a | n/a | PRJNA947942 |
| LineA_HaploidMale_6 | Nextera Flex | A | C18 | Haploid Male | 25.3 | n/a | n/a | PRJNA947942 |
| LineB_HaploidMale_4 | Nextera Flex | B | STC1 | Haploid Male | 31.0 | n/a | n/a | PRJNA947942 |
| LineB_HaploidMale_5 | Nextera Flex | B | STC12 | Haploid Male | 32.8 | n/a | n/a | PRJNA947942 |
| LinkedReadPedigree2_Sister_1 | TELL-Seq | B | STC6 – dsRed | Diploid Female | 74.7 | 0.61 | 28169 | PRJNA947942 |
| LinkedReadPedigree2_Sister_2 | TELL-Seq | B | STC6 – dsRed | Diploid Female | 196.7 | 0.63 | 32090 | PRJNA947942 |
| LinkedReadPedigree1_Mother | TELL-Seq | B | STC6 – dsRed | Diploid Female | 221.5 | 0.64 | 32510 | PRJNA947942 |
| LinkedReadPedigree1_Daughter | TELL-Seq | B | STC6 – dsRed | Diploid Female | 90 | 0.62 | 29153 | PRJNA947942 |
| LinkedReadPedigree1_Daughter_TechnicalReplicateUsedAsNegativeControl | TELL-Seq | B | STC6 – dsRed | Diploid Female | 70.1 | 0.60 | 25555 | PRJNA947942 |
| LinkedReadPedigree3_Daughter_1 | TELL-Seq | B | STC6 – dsRed | Diploid Female | 73.6 | 0.60 | 27332 | PRJNA947942 |
| LinkedReadPedigree3_Mother | TELL-Seq | B | STC6 – dsRed | Diploid Female | 50.1 | 0.53 | 15665 | PRJNA947942 |
| LinkedReadPedigree3_Daughter_2 | TELL-Seq | B | STC6 – dsRed | Diploid Female | 100.8 | 0.62 | 28841 | PRJNA947942 |
| LinkedReadPedigree4_Mother | TELL-Seq | B | STC6 – dsRed | Diploid Female | 92.4 | 0.61 | 28875 | PRJNA947942 |
| LinkedReadPedigree4_Daughter_1 | TELL-Seq | B | STC6 – dsRed | Diploid Female | 91.1 | 0.61 | 27742 | PRJNA947942 |
| LinkedReadPedigree4_Daughter_2 | TELL-Seq | B | STC6 – dsRed | Diploid Female | 81 | 0.61 | 28477 | PRJNA947942 |
| LinkedReadPedigree5_Sister_1 | TELL-Seq | B | STC6 – dsRed | Diploid Female | 75.5 | 0.61 | 27487 | PRJNA947942 |
| LinkedReadPedigree5_Sister_2 | TELL-Seq | B | STC6 – dsRed | Diploid Female | 70 | 0.59 | 25356 | PRJNA947942 |
| LinkedReadPedigree6_Sister_1 | TELL-Seq | B | STC6 – dsRed | Diploid Female | 77.6 | 0.61 | 29383 | PRJNA947942 |
| LinkedReadPedigree6_Sister_2 | TELL-Seq | B | STC6 – dsRed | Diploid Female | 69.3 | 0.59 | 25043 | PRJNA947942 |
| LinkedReadPedigree7_Sister_1 | TELL-Seq | B | STC6 - dsRed | Diploid Female | 91.9 | 0.62 | 29219 | PRJNA947942 |
| LinkedReadPedigree7_Sister_2 | TELL-Seq | B | STC6 - dsRed | Diploid Female | 82.3 | 0.62 | 29001 | PRJNA947942 |

**Supplementary Table 3.** Genome-wide heterozygosity data for all individuals from known pedigrees.

| Sample name | Library prep method | Number of windows containing a heterozygous SNP | Total number of non-overlapping 10kb windows | Proportion of windows containing a heterozygous SNP |
| --- | --- | --- | --- | --- |
| ShortReadPedigree1_Mother | Nextera Flex | 19897 | 21784 | 0.91337679 |
| ShortReadPedigree1_Daughter | Nextera Flex | 19897 | 21784 | 0.91337679 |
| ShortReadPedigree2_Mother | Nextera Flex | 19898 | 21784 | 0.913422696 |
| ShortReadPedigree2_Daughter | Nextera Flex | 19898 | 21784 | 0.913422696 |
| LinkedReadPedigree3_Daughter_1 | TELL-Seq | 19870 | 21784 | 0.912137349 |
| LinkedReadPedigree3_Mother | TELL-Seq | 19870 | 21784 | 0.912137349 |
| LinkedReadPedigree3_Daughter_2 | TELL-Seq | 19870 | 21784 | 0.912137349 |
| LinkedReadPedigree4_Mother | TELL-Seq | 19871 | 21784 | 0.912183254 |
| LinkedReadPedigree4_Daughter_1 | TELL-Seq | 19871 | 21784 | 0.912183254 |
| LinkedReadPedigree4_Daughter_2 | TELL-Seq | 19871 | 21784 | 0.912183254 |
| LinkedReadPedigree5_Sister_1 | TELL-Seq | 19871 | 21784 | 0.912183254 |
| LinkedReadPedigree5_Sister_2 | TELL-Seq | 19871 | 21784 | 0.912183254 |
| LinkedReadPedigree6_Sister_1 | TELL-Seq | 19871 | 21784 | 0.912183254 |
| LinkedReadPedigree6_Sister_2 | TELL-Seq | 19871 | 21784 | 0.912183254 |
| LinkedReadPedigree7_Sister_1 | TELL-Seq | 19850 | 21784 | 0.911219243 |
| LinkedReadPedigree7_Sister_2 | TELL-Seq | 19850 | 21784 | 0.911219243 |
| LinkedReadPedigree2_Sister_1 | TELL-Seq | 19871 | 21784 | 0.912183254 |
| LinkedReadPedigree2_Sister_2 | TELL-Seq | 19871 | 21784 | 0.912183254 |
| LinkedReadPedigree1_Mother | TELL-Seq | 19871 | 21784 | 0.912183254 |
| LinkedReadPedigree1_Daughter | TELL-Seq | 19871 | 21784 | 0.912183254 |
| LinkedReadPedigree1_Daughter_Technical ReplicateUsedAsNegativeControl | TELL-Seq | 19871 | 21784 | 0.912183254 |

**Supplementary Table 4.** All SNPs that lost or gained heterozygosity in known-pedigree pairs. LOH: loss of heterozygosity.

| Position on chromosomal scaffold | Sample 1 genotype | Sample 2 genotype | Multi-SNP event? | Pair of samples that the LOH was detected between | Number of intervening meioses between samples |
| --- | --- | --- | --- | --- | --- |
| Chr11:2981030 | C/T | T/T | No | SRPed1_Mother, SRPed1_Daughter | 1 |
| Chr3:2919542 | A/A | A/C | No | SRPed2_Mother, SRPed2_Daughter | 1 |
| Chr1:2880482 | C/T | C/C | No | SRPed1_Mother, SRPed2_Mother | 4-20 |
| Chr1:3541919 | T/T | T/C | No | SRPed1_Mother, SRPed2_Mother | 4-20 |
| Chr2:8055143 | T/C | C/C | Yes, with Chr2:8055262 and Chr2:8055673 | SRPed1_Mother, SRPed2_Mother | 4-20 |
| Chr2:8055262 | G/T | T/T | Yes, with Chr2:8055143 and Chr2:8055673 | SRPed1_Mother, SRPed2_Mother | 4-20 |
| Chr2:8055673 | T/C | C/C | Yes, with Chr2:8055143 and Chr2:8055262 | SRPed1_Mother, SRPed2_Mother | 4-20 |
| Chr3:12401182 | T/T | C/T | No | SRPed1_Mother, SRPed2_Mother | 4-20 |
| Chr5:2998673 | T/T | C/T | No | SRPed1_Mother, SRPed2_Mother | 4-20 |
| Chr8:8767770 | C/T | C/C | No | SRPed1_Mother, SRPed2_Mother | 4-20 |
| Chr9:5958719 | G/G | G/A | No | SRPed1_Mother, SRPed2_Mother | 4-20 |
| Chr9:6758788 | G/G | G/T | No | SRPed1_Mother, SRPed2_Mother | 4-20 |
| Chr10:4817986 | T/T | C/T | Yes, with Chr10:4818010 | SRPed1_Mother, SRPed2_Mother | 4-20 |
| Chr10:4818010 | T/T | C/T | Yes, with Chr10:4817986 | SRPed1_Mother, SRPed2_Mother | 4-20 |
| Chr11:10844572 | A/A | G/A | No | SRPed1_Mother,<br>SRPed2_Mother | 4-20 |
| Chr12:3309672 | T/C | T/T | No | SRPed1_Mother,<br>SRPed2_Mother | 4-20 |
| Chr14:6563773 | C/C | C/T | Yes, with<br>Chr14:6563775 | SRPed1_Mother,<br>SRPed2_Mother | 4-20 |
| Chr14:6563775 | A/A | A/G | Yes, with<br>Chr14:6563773 | SRPed1_Mother,<br>SRPed2_Mother | 4-20 |
| Chr14:6810817 | G/A | G/G | No | SRPed1_Mother,<br>SRPed2_Mother | 4-20 |
| Chr14:7159021 | A/G | A/A | No | SRPed1_Mother,<br>SRPed2_Mother | 4-20 |
| chr1:2636278 | G/A | G/G | No | LRPed2_Sister_1,<br>LRPed2_Sister_2 | 2 |
| chr11:4229141 | C/T | T/T | No | LRPed2_Sister_1,<br>LRPed2_Sister_2 | 2 |
| chr1:9586051 | C/C | C/A | No | LRPed3_Daughter_1,<br>LRPed3_Mother | 1 |
| chr1:21528656 | T/C | C/C | No | LRPed3_Daughter_1,<br>LRPed3_Mother | 1 |
| chr2:549944 | A/A | G/A | No | LRPed3_Daughter_1,<br>LRPed3_Mother | 1 |
| chr5:14571357 | C/C | C/T | No | LRPed3_Daughter_1,<br>LRPed3_Mother | 1 |
| chr1:3432379 | C/T | C/C | No | LRPed3_Mother,<br>LRPed3_Daughter_2 | 1 |
| chr3:9391605 | C/C | A/C | No | LRPed3_Mother,<br>LRPed3_Daughter_2 | 1 |
| chr8:10166826 | G/A | G/G | No | LRPed3_Mother,<br>LRPed3_Daughter_2 | 1 |
| chr4:17378017 | A/G | G/G | No | LRPed4_Mother,<br>LRPed4_Daughter_1 | 1 |
| chr5:12122624 | C/C | C/T | No | LRPed4_Mother,<br>LRPed4_Daughter_1 | 1 |
| chr9:9400932 | A/T | A/A | No | LRPed4_Mother,<br>LRPed4_Daughter_1 | 1 |
| chr4:4783832 | T/G | T/T | No | LRPed5_Sister_1,<br>LRPed5_Sister_2 | 2 |
| chr5:15866169 | T/C | C/C | No | LRPed6_Sister_1,<br>LRPed6_Sister_2 | 2 |
| chr12:2082500 | C/C | C/T | No | LRPed6_Sister_1,<br>LRPed6_Sister_2 | 2 |

**Supplementary Table 5.** All crossovers detected in comparisons of phased genomes. Each row reports one observed crossover, where the relative phasing of samples changed from SNP1 to SNP2. If loss of heterozygosity occurred in the vicinity of the crossover, the identity of the SNPs at which it occurred, and the minimum and maximum tract lengths of this loss of heterozygosity event are given. LOH: loss of heterozygosity.

| Chromosome | SNP1 | SNP2 | Associated SNPs that bear LOH | Min. length of LOH tract (bp) | Max. length of LOH tract (bp) | Pair of samples that the crossover was detected between | Pair type |
| --- | --- | --- | --- | --- | --- | --- | --- |
| Chr1 | 3144729 | 3146898 | None | n/a | n/a | LRPed4_Mother, LRPed4_Daughter_2 | Mother-Daughter |
| Chr1 | 3431086 | 3433123 | Chr1:3432379 | 1 | 2036 | LRPed3_Mother, LRPed3_Daughter_2 | Mother-Daughter |
| Chr2 | 9547071 | 9549155 | None | n/a | n/a | LRPed4_Mother, LRPed4_Daughter_2 | Mother-Daughter |
| Chr3 | 11210015 | 11210761 | None | n/a | n/a | LRPed1_Mother, LRPed1_Daughter | Mother-Daughter |
| Chr5 | 15025316 | 15028015 | None | n/a | n/a | LRPed4_Mother, LRPed4_Daughter_1 | Mother-Daughter |
| Chr6 | 2474943 | 2476664 | None | n/a | n/a | LRPed3_Mother, LRPed3_Daughter_1 | Mother-Daughter |
| Chr6 | 2542519 | 2546612 | None | n/a | n/a | LRPed4_Mother, LRPed4_Daughter_2 | Mother-Daughter |
| Chr9 | 9398135 | 9402024 | Chr9:9400932 | 1 | 3888 | LRPed4_Mother, LRPed4_Daughter_1 | Mother-Daughter |
| Chr11 | 4190454 | 4191627 | None | n/a | n/a | LRPed4_Mother, LRPed4_Daughter_1 | Mother-Daughter |
| Chr11 | 6108203 | 6115529 | None | n/a | n/a | LRPed1_Mother, LRPed1_Daughter | Mother-Daughter |
| Chr12 | 4977421 | 4978079 | None | n/a | n/a | LRPed3_Mother, LRPed3_Daughter_1 | Mother-Daughter |
| Chr14 | 6749385 | 6749925 | None | n/a | n/a | LRPed1_Mother, LRPed1_Daughter | Mother-Daughter |
| Chr1 | 2634620 | 2636619 | Chr1:2636278 | 1 | 1998 | LRPed2_Sister_1,<br>LRPed2_Sister_2 | Sister-Sister |
| Chr1 | 3229322 | 3231706 | None | n/a | n/a | LRPed6_Sister_1,<br>LRPed6_Sister_2 | Sister-Sister |
| Chr1 | 5807036 | 5809174 | None | n/a | n/a | LRPed5_Sister_1,<br>LRPed5_Sister_2 | Sister-Sister |
| Chr2 | 7327271 | 7330336 | None | n/a | n/a | LRPed7_Sister_1,<br>LRPed7_Sister_2 | Sister-Sister |
| Chr3 | 142174 | 144749 | None | n/a | n/a | LRPed7_Sister_1,<br>LRPed7_Sister_2 | Sister-Sister |
| Chr3 | 2370477 | 2372682 | None | n/a | n/a | LRPed6_Sister_1,<br>LRPed6_Sister_2 | Sister-Sister |
| Chr3 | 12979809 | 12982121 | None | n/a | n/a | LRPed2_Sister_1,<br>LRPed2_Sister_2 | Sister-Sister |
| Chr4 | 4783400 | 4784862 | Chr4:4783832 | 1 | 1461 | LRPed5_Sister_1,<br>LRPed5_Sister_2 | Sister-Sister |
| Chr4 | 14174116 | 14177470 | None | n/a | n/a | LRPed7_Sister_1,<br>LRPed7_Sister_2 | Sister-Sister |
| Chr5 | 15865540 | 15868891 | Chr5:15866169 | 1 | 3350 | LRPed6_Sister_1,<br>LRPed6_Sister_2 | Sister-Sister |
| Chr7 | 1046985 | 1049060 | None | n/a | n/a | LRPed2_Sister_1,<br>LRPed2_Sister_2 | Sister-Sister |
| Chr7 | 2995655 | 3000576 | None | n/a | n/a | LRPed6_Sister_1,<br>LRPed6_Sister_2 | Sister-Sister |
| Chr7 | 10767949 | 10772010 | None | n/a | n/a | LRPed2_Sister_1,<br>LRPed2_Sister_2 | Sister-Sister |
| Chr8 | 9557056 | 9559006 | None | n/a | n/a | LRPed6_Sister_1,<br>LRPed6_Sister_2 | Sister-Sister |
| Chr9 | 15375104 | 15376819 | None | n/a | n/a | LRPed6_Sister_1,<br>LRPed6_Sister_2 | Sister-Sister |
| Chr10 | 4107179 | 4110860 | None | n/a | n/a | LRPed2_Sister_1,<br>LRPed2_Sister_2 | Sister-Sister |
| Chr11 | 3295416 | 3301382 | None | n/a | n/a | LRPed7_Sister_1,<br>LRPed7_Sister_2 | Sister-Sister |
| Chr11 | 4228467 | 4230142 | None | n/a | n/a | LRPed2_Sister_1,<br>LRPed2_Sister_2 | Sister-Sister |
| Chr11 | 13111362 | 13113378 | None | n/a | n/a | LRPed5_Sister_1,<br>LRPed5_Sister_2 | Sister-Sister |
| Chr1 | 2505258 | 2512078 | None | n/a | n/a | See Supplementary<br>Data 4 | Unknown pedigree |
| Chr1 | 2580080 | 2581562 | None | n/a | n/a | See Supplementary Data 4 | Unknown pedigree |
| Chr1 | 3485060 | 3486472 | None | n/a | n/a | See Supplementary Data 4 | Unknown pedigree |
| Chr1 | 3585374 | 3587447 | None | n/a | n/a | See Supplementary Data 4 | Unknown pedigree |
| Chr1 | 4230737 | 4234105 | None | n/a | n/a | See Supplementary Data 4 | Unknown pedigree |
| Chr1 | 5254671 | 5257204 | None | n/a | n/a | See Supplementary Data 4 | Unknown pedigree |
| Chr1 | 5929741 | 5932776 | None | n/a | n/a | See Supplementary Data 4 | Unknown pedigree |
| Chr1 | 15392918 | 15394244 | None | n/a | n/a | See Supplementary Data 4 | Unknown pedigree |
| Chr1 | 20094169 | 20095268 | None | n/a | n/a | See Supplementary Data 4 | Unknown pedigree |
| Chr1 | 20105229 | 20108062 | None | n/a | n/a | See Supplementary Data 4 | Unknown pedigree |
| Chr2 | 8687309 | 8689810 | None | n/a | n/a | See Supplementary Data 4 | Unknown pedigree |
| Chr2 | 9466957 | 9468997 | Chr2:9467184,<br>Chr2:9467479,<br>Chr2:9467650 | 467 | 2039 | See Supplementary Data 4 | Unknown pedigree |
| Chr2 | 13288306 | 13290678 | None | n/a | n/a | See Supplementary Data 4 | Unknown pedigree |
| Chr2 | 16649229 | 16650349 | None | n/a | n/a | See Supplementary Data 4 | Unknown pedigree |
| Chr2 | 17408232 | 17409489 | None | n/a | n/a | See Supplementary Data 4 | Unknown pedigree |
| Chr2 | 17741774 | 17743684 | None | n/a | n/a | See Supplementary Data 4 | Unknown pedigree |
| Chr3 | 2825664 | 2830772 | None | n/a | n/a | See Supplementary Data 4 | Unknown pedigree |
| Chr3 | 3450629 | 3452920 | None | n/a | n/a | See Supplementary Data 4 | Unknown pedigree |
| Chr3 | 10589952 | 10591127 | None | n/a | n/a | See Supplementary Data 4 | Unknown pedigree |
| Chr3 | 11678342 | 11679855 | None | n/a | n/a | See Supplementary Data 4 | Unknown pedigree |
| Chr3 | 13079124 | 13083881 | None | n/a | n/a | See Supplementary Data 4 | Unknown pedigree |
| Chr3 | 14734965 | 14740935 | None | n/a | n/a | See Supplementary Data 4 | Unknown pedigree |
| Chr3 | 15620239 | 15622094 | None | n/a | n/a | See Supplementary Data 4 | Unknown pedigree |
| Chr4 | 1799347 | 1800310 | None | n/a | n/a | See Supplementary Data 4 | Unknown pedigree |
| Chr4 | 4764390 | 4765375 | None | n/a | n/a | See Supplementary Data 4 | Unknown pedigree |
| Chr4 | 4961344 | 4963545 | None | n/a | n/a | See Supplementary Data 4 | Unknown pedigree |
| Chr4 | 6493439 | 6500497 | None | n/a | n/a | See Supplementary Data 4 | Unknown pedigree |
| Chr4 | 6540198 | 6544783 | None | n/a | n/a | See Supplementary Data 4 | Unknown pedigree |
| Chr4 | 6902345 | 6907619 | None | n/a | n/a | See Supplementary Data 4 | Unknown pedigree |
| Chr4 | 12787752 | 12788448 | None | n/a | n/a | See Supplementary Data 4 | Unknown pedigree |
| Chr4 | 12893926 | 12895898 | None | n/a | n/a | See Supplementary Data 4 | Unknown pedigree |
| Chr4 | 14465460 | 14469886 | None | n/a | n/a | See Supplementary Data 4 | Unknown pedigree |
| Chr4 | 14799061 | 14801738 | None | n/a | n/a | See Supplementary Data 4 | Unknown pedigree |
| Chr5 | 1683004 | 1684622 | None | n/a | n/a | See Supplementary Data 4 | Unknown pedigree |
| Chr5 | 1742388 | 1746659 | None | n/a | n/a | See Supplementary Data 4 | Unknown pedigree |
| Chr5 | 2203037 | 2205258 | None | n/a | n/a | See Supplementary Data 4 | Unknown pedigree |
| Chr5 | 4121926 | 4124422 | None | n/a | n/a | See Supplementary Data 4 | Unknown pedigree |
| Chr5 | 4676692 | 4677079 | Chr5:4676805 | 1 | 386 | See Supplementary Data 4 | Unknown pedigree |
| Chr5 | 5024162 | 5026314 | None | n/a | n/a | See Supplementary Data 4 | Unknown pedigree |
| Chr5 | 6364075 | 6365062 | None | n/a | n/a | See Supplementary Data 4 | Unknown pedigree |
| Chr5 | 14764564 | 14765973 | None | n/a | n/a | See Supplementary Data 4 | Unknown pedigree |
| Chr5 | 16169417 | 16172134 | None | n/a | n/a | See Supplementary Data 4 | Unknown pedigree |
| Chr5 | 16487822 | 16492100 | Chr5:16488439 | 1 | 4277 | See Supplementary Data 4 | Unknown pedigree |
| Chr6 | 1293855 | 1295378 | None | n/a | n/a | See Supplementary Data 4 | Unknown pedigree |
| Chr6 | 1477808 | 1479095 | None | n/a | n/a | See Supplementary Data 4 | Unknown pedigree |
| Chr6 | 1665900 | 1669463 | Chr6:1668578 | 1 | 3562 | See Supplementary Data 4 | Unknown pedigree |
| Chr6 | 1679067 | 1682125 | None | n/a | n/a | See Supplementary Data 4 | Unknown pedigree |
| Chr6 | 2491691 | 2493148 | None | n/a | n/a | See Supplementary Data 4 | Unknown pedigree |
| Chr6 | 3383363 | 3387817 | None | n/a | n/a | See Supplementary Data 4 | Unknown pedigree |
| Chr6 | 3868688 | 3869821 | None | n/a | n/a | See Supplementary Data 4 | Unknown pedigree |
| Chr6 | 4384294 | 4386828 | Chr6:4386429 | 1 | 2533 | See Supplementary Data 4 | Unknown pedigree |
| Chr6 | 4753729 | 4757462 | None | n/a | n/a | See Supplementary Data 4 | Unknown pedigree |
| Chr6 | 4787560 | 4788227 | None | n/a | n/a | See Supplementary Data 4 | Unknown pedigree |
| Chr6 | 10459863 | 10464260 | None | n/a | n/a | See Supplementary Data 4 | Unknown pedigree |
| Chr6 | 12404559 | 12406688 | None | n/a | n/a | See Supplementary Data 4 | Unknown pedigree |
| Chr6 | 13842670 | 13846879 | None | n/a | n/a | See Supplementary Data 4 | Unknown pedigree |
| Chr7 | 1375278 | 1376502 | None | n/a | n/a | See Supplementary Data 4 | Unknown pedigree |
| Chr7 | 1576858 | 1583933 | None | n/a | n/a | See Supplementary Data 4 | Unknown pedigree |
| Chr7 | 1595992 | 1602423 | None | n/a | n/a | See Supplementary Data 4 | Unknown pedigree |
| Chr7 | 1672444 | 1678644 | None | n/a | n/a | See Supplementary Data 4 | Unknown pedigree |
| Chr7 | 1682957 | 1686147 | None | n/a | n/a | See Supplementary Data 4 | Unknown pedigree |
| Chr7 | 1842366 | 1842892 | None | n/a | n/a | See Supplementary Data 4 | Unknown pedigree |
| Chr7 | 1893203 | 1893501 | None | n/a | n/a | See Supplementary Data 4 | Unknown pedigree |
| Chr7 | 3491438 | 3494048 | None | n/a | n/a | See Supplementary Data 4 | Unknown pedigree |
| Chr7 | 4221344 | 4223148 | None | n/a | n/a | See Supplementary Data 4 | Unknown pedigree |
| Chr7 | 16350423 | 16353153 | None | n/a | n/a | See Supplementary Data 4 | Unknown pedigree |
| Chr8 | 4835591 | 4840348 | None | n/a | n/a | See Supplementary Data 4 | Unknown pedigree |
| Chr8 | 7664695 | 7665534 | None | n/a | n/a | See Supplementary Data 4 | Unknown pedigree |
| Chr8 | 9481327 | 9484026 | None | n/a | n/a | See Supplementary Data 4 | Unknown pedigree |
| Chr8 | 10052497 | 10055615 | None | n/a | n/a | See Supplementary Data 4 | Unknown pedigree |
| Chr8 | 10058727 | 10064250 | None | n/a | n/a | See Supplementary Data 4 | Unknown pedigree |
| Chr8 | 10067490 | 10072718 | None | n/a | n/a | See Supplementary Data 4 | Unknown pedigree |
| Chr8 | 10210998 | 10211797 | None | n/a | n/a | See Supplementary Data 4 | Unknown pedigree |
| Chr8 | 12837072 | 12839214 | None | n/a | n/a | See Supplementary Data 4 | Unknown pedigree |
| Chr8 | 15380104 | 15384110 | Chr8:15382125 | 1 | 4005 | See Supplementary Data 4 | Unknown pedigree |
| Chr9 | 459786 | 464426 | None | n/a | n/a | See Supplementary Data 4 | Unknown pedigree |
| Chr9 | 462909 | 464426 | None | n/a | n/a | See Supplementary Data 4 | Unknown pedigree |
| Chr9 | 9876319 | 9880695 | None | n/a | n/a | See Supplementary Data 4 | Unknown pedigree |
| Chr9 | 10116851 | 10119633 | None | n/a | n/a | See Supplementary Data 4 | Unknown pedigree |
| Chr9 | 10228908 | 10233305 | Chr9:10231668 | 1 | 4396 | See Supplementary Data 4 | Unknown pedigree |
| Chr9 | 12093888 | 12094936 | Chr9:12094339 | 1 | 1047 | See Supplementary Data 4 | Unknown pedigree |
| Chr9 | 12455642 | 12457378 | Chr9:1245662<br>3 | 1 | 1735 | See Supplementary<br>Data 4 | Unknown<br>pedigree |
| Chr9 | 12859913 | 12864936 | None | n/a | n/a | See Supplementary<br>Data 4 | Unknown<br>pedigree |
| Chr9 | 14484131 | 14487391 | None | n/a | n/a | See Supplementary<br>Data 4 | Unknown<br>pedigree |
| Chr9 | 14599644 | 14602284 | None | n/a | n/a | See Supplementary<br>Data 4 | Unknown<br>pedigree |
| Chr10 | 7212895 | 7213764 | None | n/a | n/a | See Supplementary<br>Data 4 | Unknown<br>pedigree |
| Chr10 | 7819982 | 7823644 | None | n/a | n/a | See Supplementary<br>Data 4 | Unknown<br>pedigree |
| Chr11 | 2704615 | 2705595 | None | n/a | n/a | See Supplementary<br>Data 4 | Unknown<br>pedigree |
| Chr11 | 3764960 | 3771086 | None | n/a | n/a | See Supplementary<br>Data 4 | Unknown<br>pedigree |
| Chr11 | 4011872 | 4014721 | None | n/a | n/a | See Supplementary<br>Data 4 | Unknown<br>pedigree |
| Chr11 | 4182567 | 4185672 | None | n/a | n/a | See Supplementary<br>Data 4 | Unknown<br>pedigree |
| Chr11 | 4242592 | 4245310 | None | n/a | n/a | See Supplementary<br>Data 4 | Unknown<br>pedigree |
| Chr11 | 4468337 | 4470202 | None | n/a | n/a | See Supplementary<br>Data 4 | Unknown<br>pedigree |
| Chr11 | 5294618 | 5299679 | Chr11:529543<br>3 | 1 | 5060 | See Supplementary<br>Data 4 | Unknown<br>pedigree |
| Chr11 | 5608575 | 5610360 | None | n/a | n/a | See Supplementary<br>Data 4 | Unknown<br>pedigree |
| Chr11 | 5866085 | 5869277 | None | n/a | n/a | See Supplementary<br>Data 4 | Unknown<br>pedigree |
| Chr11 | 9299318 | 9302235 | None | n/a | n/a | See Supplementary<br>Data 4 | Unknown<br>pedigree |
| Chr11 | 13423045 | 13427194 | Chr11:134242<br>68,<br>Chr11:134243<br>30 | 63 | 4148 | See Supplementary<br>Data 4 | Unknown<br>pedigree |
| Chr11 | 13568583 | 13572052 | None | n/a | n/a | See Supplementary<br>Data 4 | Unknown<br>pedigree |
| Chr12 | 3259779 | 3261408 | None | n/a | n/a | See Supplementary<br>Data 4 | Unknown<br>pedigree |
| Chr12 | 3376563 | 3379548 | None | n/a | n/a | See Supplementary Data 4 | Unknown pedigree |
| Chr12 | 4211216 | 4213893 | None | n/a | n/a | See Supplementary Data 4 | Unknown pedigree |
| Chr12 | 4716727 | 4719898 | None | n/a | n/a | See Supplementary Data 4 | Unknown pedigree |
| Chr12 | 4988162 | 4992987 | None | n/a | n/a | See Supplementary Data 4 | Unknown pedigree |
| Chr12 | 6108980 | 6111583 | None | n/a | n/a | See Supplementary Data 4 | Unknown pedigree |
| Chr12 | 6964881 | 6967149 | None | n/a | n/a | See Supplementary Data 4 | Unknown pedigree |
| Chr12 | 7381401 | 7384669 | None | n/a | n/a | See Supplementary Data 4 | Unknown pedigree |
| Chr12 | 7685985 | 7688635 | Chr12:768774<br>7 | 1 | 2649 | See Supplementary Data 4 | Unknown pedigree |
| Chr14 | 3796921 | 3799968 | None | n/a | n/a | See Supplementary Data 4 | Unknown pedigree |
| Chr14 | 6792999 | 6795016 | None | n/a | n/a | See Supplementary Data 4 | Unknown pedigree |
| Chr14 | 6824360 | 6828482 | None | n/a | n/a | See Supplementary Data 4 | Unknown pedigree |
| Chr14 | 7129631 | 7132302 | None | n/a | n/a | See Supplementary Data 4 | Unknown pedigree |
| Chr14 | 7593751 | 7602804 | 74 SNPs from<br>Chr14:760280<br>4-7804865 | 202061 | 211114 | See Supplementary Data 4 | Unknown pedigree |

**Supplementary Video 1**

Z-stack images of a DAPI-stained metaphase I nucleus taken at intervals of 0.17 µm. This oocyte was incompletely squashed, and so the alignment of chiasmate chromosomes during metaphase can be clearly seen.

**Supplementary Data 1**

Haplotype blocks and loss of heterozygosity tracts produced by crossing over and gene conversion among individuals from the same colony, including events observed among haploid male genomes and inferred via loss of heterozygosity among diploid female genomes. For haplotype blocks among haploids, the values indicate phasing relative to the first individual— values of “0” indicate that this individual has the same allele as the first individual for this haplotype, whereas values of “1” indicate that this individual possesses the alternate allele. For losses of heterozygosity among diploids, values of “1” mean that an individual possesses that loss of heterozygosity tract, whereas values of “0” indicate that an individual is heterozygous for that tract. For crossovers inferred from loss of heterozygosity tracts, crossovers are counted at the beginning or end of a loss of heterozygosity tract if it was not also the first or last SNP on a contig.

**Supplementary Data 2**

Haplotype blocks and loss of heterozygosity tracts produced by crossing over and gene conversion among individuals from different colonies, including events observed among haploid male genomes and inferred via loss of heterozygosity among diploid female genomes. For haplotype blocks among haploids, the values indicate phasing relative to the first individual— values of “0” indicate that this individual has the same allele as the first individual for this haplotype, whereas values of “1” indicate that this individual possesses the alternate allele. For losses of heterozygosity among diploids, values of “1” mean that an individual possesses that loss of heterozygosity tract, whereas values of “0” indicate that an individual is heterozygous for that tract.

**Supplementary Data 3**

Loss of heterozygosity tracts produced by crossing over and gene conversion among individuals from different known pedigrees from linked-read sequencing experiments. Values of “1” mean that an individual possesses that loss of heterozygosity tract, whereas values of “0” indicate that an individual is heterozygous for that tract.

**Supplementary Data 4**

Crossovers detected in all pairwise comparisons between representatives from each linked-read sequencing pedigree. For each crossover, we label the two different haplotypes for the two focal SNPs as 0 and 1.

